# Diverse mating phenotypes impact the spread of *wtf* meiotic drivers in *S. pombe*

**DOI:** 10.1101/2021.05.28.446231

**Authors:** José Fabricio López Hernández, Rachel M. Helston, Jeffrey J. Lange, R. Blake Billmyre, Samantha H. Schaffner, Michael T. Eickbush, Scott McCroskey, Sarah E. Zanders

## Abstract

Meiotic drivers are genetic loci that break Mendel’s law of segregation to be transmitted into more than half of the offspring produced by a heterozygote. The success of a driver relies on outcrossing because drivers gain their advantage in heterozygotes. It is, therefore, curious that *Schizosaccharomyces pombe*, a species reported to rarely outcross, harbors many meiotic drivers. To address this paradox, we measured mating phenotypes in *S. pombe* natural isolates. We found that the propensity to inbreed varies between natural isolates and can be affected both by cell density and by the available sexual partners. Additionally, we found that the observed level of inbreeding slows, but does not prevent, the spread of a *wtf* meiotic driver in the absence of additional fitness costs. These analyses reveal parameters critical to understanding the evolution of *S. pombe* and help explain the success of meiotic drivers in this species.

## Introduction

Mating behaviours have long been a focus of art, literature, and formal scientific inquiry. This interest stems, in part, from the remarkable importance of mate choice on the evolution of species. Outcrossing and inbreeding represent distinct mating strategies that both have potential evolutionary benefits and costs (Glemin et al. 2019; Muller 1932; Otto and Lenormand 2002). For example, preferential outcrossing can facilitate the spread of adaptive traits in a population, but can also promote the spread of deleterious selfish genes (Crow 1988; Hurst and Werren 2001; McDonald et al. 2016; Zeyl et al. 1996).

Meiotic drivers represent one type of selfish genetic element that relies on outcrossing to persist and spread in a population (Lindholm et al. 2016; Novitski 1957). These loci can manipulate the process of gametogenesis to bias their own transmission into gametes at the expense of the rest of the genome (Burt and Trivers 2006). Meiotic drivers are often considered selfish or parasitic genes because they generally offer no fitness benefits to their hosts and are instead often deleterious or linked to deleterious alleles (Dyer et al. 2007; Higgins et al. 2018; Klein et al. 1984; Rick 1966; Schimenti et al. 2005; Taylor et al. 1999; Wilkinson and Fry 2001). As inbreeding is thought to inhibit the spread of selfish genes like drivers, drivers are predicted to be unsuccessful in species that rarely outcross (Hurst and Werren 2001). This assumption appears to be challenged in the fission yeast *Schizosaccharomyces pombe,* which is thought to rarely outcross, yet hosts multiple meiotic drivers (Bravo Nunez et al. 2020b; Eickbush et al. 2019; Farlow et al. 2015; Hu et al. 2017; Nuckolls et al. 2017; Tusso et al. 2019; Zanders et al. 2014).

In the wild, a minority of *S. pombe* strains are heterothallic, meaning they have a fixed mating type (*h+* or *h-*; Gutz and Doe 1975; Nieuwenhuis and Immler 2016; Schlake 1993). Heterothallic strains must outcross to complete sexual reproduction (Egel 1977; Gutz and Doe 1975; Leupold 1949; Miyata 1981; Nieuwenhuis and Immler 2016; Osterwalder 1924; Schlake 1993). However, most isolates of *S. pombe* are homothallic, meaning that they can switch between the two mating types (*h+* and *h-*) (Eie 1987; Gutz and Doe 1975; Klar 1990; Nieuwenhuis et al. 2018; Singh and Klar 2003). Homothallism can facilitate inbreeding since mating can occur between clonal members of a population. Mating can even occur between the two sibling cells produced by a mitotic cell division (Egel 1977; Gutz and Doe 1975; Miyata 1981). In the lab, homothallic strains are thought to preferentially inbreed (Ekwall and Thon 2017; Forsburg and Rhind 2006; Merlini et al. 2013).

Population genetic analyses have provided additional support for the notion that *S. pombe* inbreeds. *S. pombe* has been estimated to outcross once every ∼800,000 generations, which is about ten-fold less frequent than predictions for the homothallic budding yeast *Saccharomyces cerevisiae* (Farlow et al. 2015). There are two major *S. pombe* lineages that diverged between 2,300 and 78,000 years ago (Tao et al. 2019; Tusso et al. 2019). Much of the sampled variation within the species represents different admixed hybrids of those two ancestral lineages resulting from an estimated 20-60 outcrossing events (Tusso et al. 2019). This low outcrossing rate could result from limited opportunities to outcross, preferential inbreeding, or a combination of the two. It is important to note, however, that the outcrossing rate estimates in *S. pombe* are likely low as pervasive meiotic drive and decreased recombination in heterozygotes suppress the genetic signatures that are used to infer outcrossing (Bravo Nunez et al. 2020b; Hu et al. 2017; Nuckolls et al. 2017; Zanders et al. 2014).

The current outcrossing rate estimates suggest that drivers would infrequently have the opportunity to act in *S. pombe*. Nonetheless, the *S. pombe* genome houses numerous meiotic drive genes from the *wtf* gene family (Eickbush et al. 2019; Hu et al. 2017; Nuckolls et al. 2017). The *wtf* drivers destroy the meiotic products (spores) that do not inherit the driver from a heterozygote. Each *wtf* drive gene encodes both a Wtf^poison^ and an Wtf^antidote^ protein that, together, execute targeted spore killing of the spores that do not inherit the *wtf* driver. In the characterized *wtf4* driver, the Wtf4^poison^ protein assembles into toxic protein aggregates that are packaged into all developing spores. The Wtf4^antidote^ protein co-assembles with Wtf4^poison^ only in the spores that inherit *wtf4* and likely neutralizes the poison by promoting its trafficking to the vacuole (Nuckolls et al. 2020). Spore killing by *wtf* drivers leads to the loss of about half of the spores and almost exclusive transmission (>90%) of the *wtf* driver from a heterozygote (Hu et al. 2017; Nuckolls et al. 2017). Despite their heavy costs in heterozygotes, the drivers are successful in that all assayed *S. pombe* isolates contain multiple *wtf* drivers and some contain ten or more predicted drivers (Bravo Nunez et al. 2020a; Eickbush et al. 2019).

In this work, we exploited the tractability of *S. pombe* to better understand how meiotic drivers could succeed. Despite limited genetic diversity amongst isolates, we observed natural variation in inbreeding propensity and other mating phenotypes. Some natural isolates preferentially inbreed in the presence of a potential outcrossing partner, whereas others mate more randomly. Additionally, we found that the level of inbreeding can be altered by cell density and affected by the available sexual partners. To explore the effects of these mating phenotypes on the spread of a *wtf* driver in a population, we used both mathematical modelling and an experimental evolution approach. We found that while the spread of a *wtf* driver could be slowed by inbreeding, the driver could still spread in the absence of linked deleterious traits. We incorporated our observations into a model in which rapid *wtf* gene evolution and occasional outcrossing facilitate the maintenance of *wtf* drivers. More broadly, this study illustrates how the success of drive systems are impacted by mating phenotypes.

## Results

### Inbreeding propensity differs amongst *S. pombe* natural isolates

Most laboratory investigations of *S. pombe* utilize cells derived from 968 (CBS1042), the first haploid isolated from French wine in 1921 by A. Osterwalder (Osterwalder 1924). In this work, we will refer to derivatives of this isolate as “*Sp*.” Homothallic *Sp* strains switch mating type in a regular pattern such that a clonal population contains an equal number of cells from each mating type (*h+* and *h-*) after relatively few cell divisions (Maki et al. 2018). When starved, haploid *S. pombe* cells of opposite mating types can mate (fuse) to form diploid zygotes, which then generally proceed directly to undergo meiosis and spore formation (sporulation). Genetics and microscopy experiments have revealed that the homothallic *Sp* haploids tend to inbreed (Bendezu and Martin 2013; Egel 1977; Leupold 1949; Merlini et al. 2016; Miyata 1981).

However, the precise level of inbreeding when *S. pombe* cells are among nonclonal sexual partners has not, to our knowledge, been formally reported for any isolate.

To quantify inbreeding in *Sp*, we first generated fluorescently tagged strains to easily observe mating via microscopy (Figure 1A). We marked strains with either GFP or mCherry (both constitutively-expressed and integrated at the *ura4* locus). We then mixed equal proportions of GFP-expressing and mCherry-expressing haploid cells and plated them on a medium (SPA) that induces mating and meiosis. We imaged the cells immediately after plating to measure the starting frequency of both parent types. We then imaged again 24-48 hours later when many cells in the population had mated to either form zygotes or fully developed spores. We inferred the genotypes (homozygous or heterozygous) of each zygote and ascus (spore sac) based on their fluorescence (Figure 1A). Homozygotes were produced by mating of two cells carrying the same fluorophore, while heterozygotes were produced by mating between a GFP-labeled and an mCherry-labeled cell (Figure 1B and C). Finally, we calculated the inbreeding coefficient (*F*) by comparing the observed frequency of heterozygotes to the frequency expected if mating was random (*F*=1-observed heterozygotes/expected heterozygotes; Figure 1A). Total inbreeding, random mating, and total outcrossing would yield inbreeding coefficients of 1, 0, and -1, respectively (Hartl and Clark 2007).

**Figure 1.**
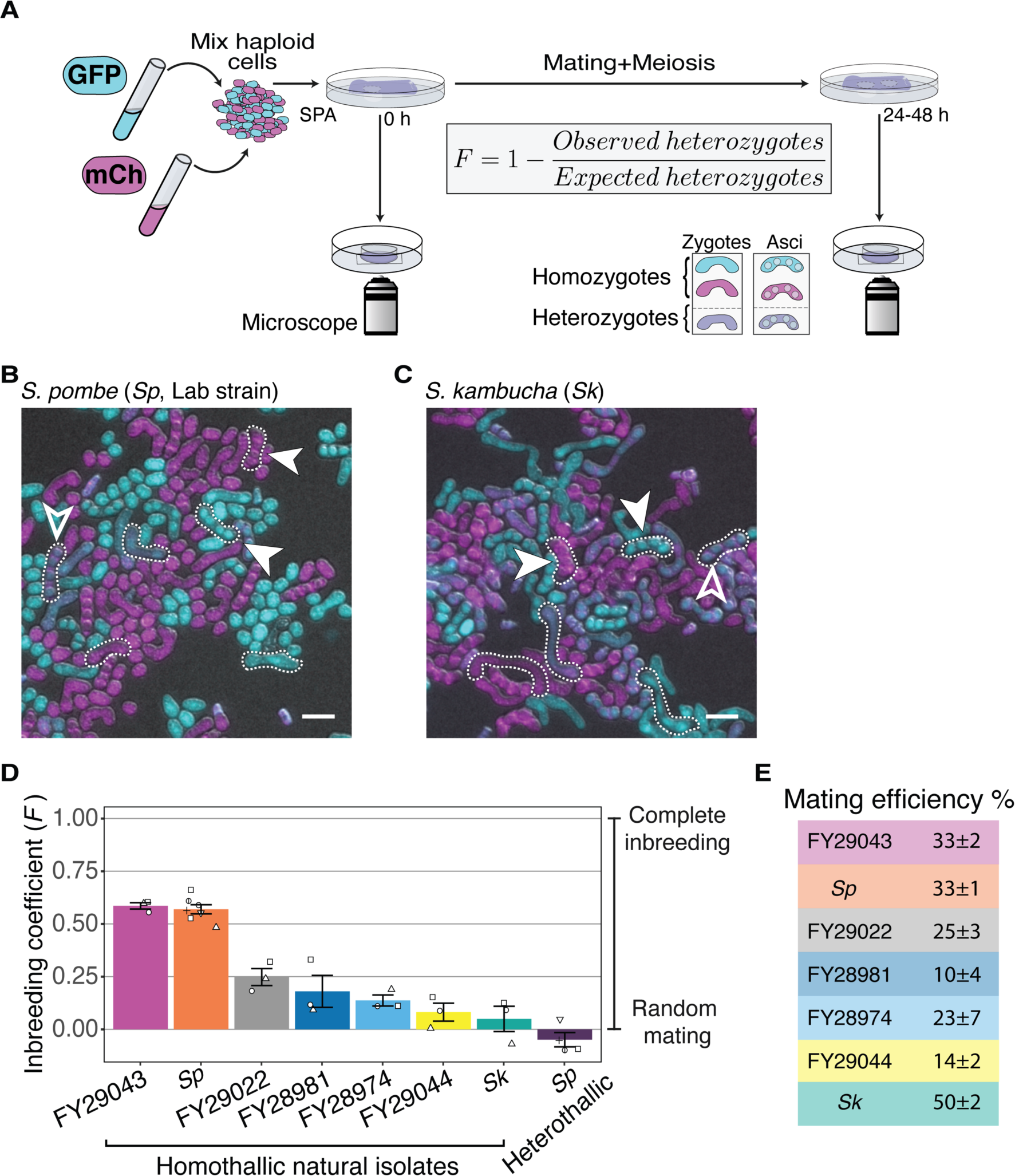
Inbreeding propensity varies between homothallic isolates of *S. pombe*. **A)** Experimental strategy to quantify inbreeding. GFP (cyan)- and mCherry (magenta)-expressing cells were mixed and placed on SPA medium that induces mating and meiosis. An agar punch from this plate was imaged to assess the initial frequencies of each haploid strain. After incubation at 25°C for at least 24 hours, another punch was imaged to determine the number of homozygous and heterozygous zygotes/asci based on their fluorescence. The inbreeding coefficient (*F*) was calculated using the formula shown. **B and C)** Representative images of the mating in *Sp* (**B**) and *Sk* (**C**) isolates after 24 hours. Filled arrowheads highlight examples of homozygous asci whereas open arrowheads highlight heterozygous asci. A few additional zygotes are also outlined with dotted lines in the images. Scale bars represent 10 µm. **D)** Inbreeding coefficient of homothallic natural isolates and complementary heterothallic (*h+,h-* mCherry and *h+,h-*^-^ *GFP*) *Sp* lab strains. At least three biological replicates per isolate are shown (open shapes). **E)** Mating efficiency of the isolates shown in **D** (%) +/- standard error from three biological replicates of each natural isolate.

In homothallic *Sp* cells, we measured an average inbreeding coefficient of 0.57 using our microscopy assay (Figure 1B and 1D). In a mixed heterothallic population (GFP-expressing and mCherry-expressing cells of both mating types), we observed random mating (*F* = -0.05), consistent with mating-type switching facilitating inbreeding (Figure 1D). To validate our microscopy results, we also assayed *Sp* cells using an orthogonal approach employing traditional genetic markers. For this analysis, we mixed haploid cells on supplemented SPA medium (SPAS) to induce them to mate and undergo meiosis. We then manually genotyped the progeny and used the fraction of recombinant progeny to calculate inbreeding coefficients (Supplemental Figure 1A). The average inbreeding coefficients measured using the genetic assay were very similar to the values we measured using the microscopy assay (0.49 for homothallic and 0.05 for heterothallic cells; Supplemental Figure 1B). Together, our results confirm previous observations of self-mating in homothallic *Sp* cells and quantify the level of inbreeding (Bendezu and Martin 2013; Egel 1977). In addition, we demonstrate that our fluorescence assay provides a powerful tool to observe and measure inbreeding.

We next extended our inbreeding analyses to other *S. pombe* natural isolates. We assayed six additional isolates, FY29043, FY29022, FY28981, FY28974, FY29044 and *S. kambucha* (*Sk*), using our fluorescence microscopy assay. We chose these isolates because they span the known diversity of *S. pombe*, are homothallic, sporulated well, were non-clumping, and we were able to transform them with the GFP and mCherry markers described above (Supplemental Table 1, Supplemental Figure 2) (Jeffares et al. 2015). We found that the inbreeding propensity varied significantly between the different natural isolates (Figure 1D). FY29043 inbred similarly to *Sp*, but other *S. pombe* isolates, including *Sk*, mated more randomly (Figure 1D). We also observed variation in mating efficiency ranging from 10% of cells mating in FY28981 to 50% of cells mating in *Sk* (Figure 1E).

Given that *S. pombe* cells are immobile, we thought that cell density could affect their propensity to inbreed. To test this, we compared the inbreeding coefficient of both homothallic *Sp* and *Sk* isolates at three different starting cell densities: our standard mating density (1X), high density (10X), and low density (0.1X). Because crowding prevented us from assaying high-density cells using our microscopy approach, we used the genetic assay for each condition. We found that inbreeding was increased in both *Sp* and *Sk* isolates when cell densities were reduced (Supplemental Figure 1C and 1D). This is likely because cells plated at low density tended to be physically distant from potential sexual partners that were not part of the same clonally growing patch of cells (Supplemental Figure 3). However, heterothallic *Sp* cells that cannot mate within a clonal patch of cells showed near random mating at all cell densities assayed (Supplemental Figure 1C). Overall, these experiments demonstrate that inbreeding propensity varies within *S. pombe* homothallic isolates and can be affected by cell density.

### Variation in mating-type switching could contribute to reduced inbreeding in *Sk*

We next used time-lapse imaging to determine the origins of the different inbreeding propensities, focusing on the *Sp* and *Sk* isolates. Previous work suggested that *Sk* cells have reduced mating-type switching efficiency, based on a Southern blot assaying the level of the DNA break (DSB) that initiates switching (Singh 2002). A mutation at the *mat-M* imprint site was proposed to be responsible for the reduced level of DSBs (Singh and Klar 2003). Since less mating-type switching could lead to less inbreeding, we decided to explore this idea using time-lapse assays (Miyata 1981). For these assays, we tracked the fate of individual homothallic founder cells plated on SPA at low density (0.25X to our standard mating density used above) to quantify how many mitotic generations occurred prior to the first mating event. When the first mating event occurred, we recorded the proportion of the cells present that mated (prior to the appearance of cells from next mitotic generation). We also recorded the relationships between the cells that did mate (i.e., sibling or non-sibling cells; Figure 2A). We did not consider cells that were born in mitotic generations past the one in which mating first occurred (Figure 2A).

**Figure 2.**
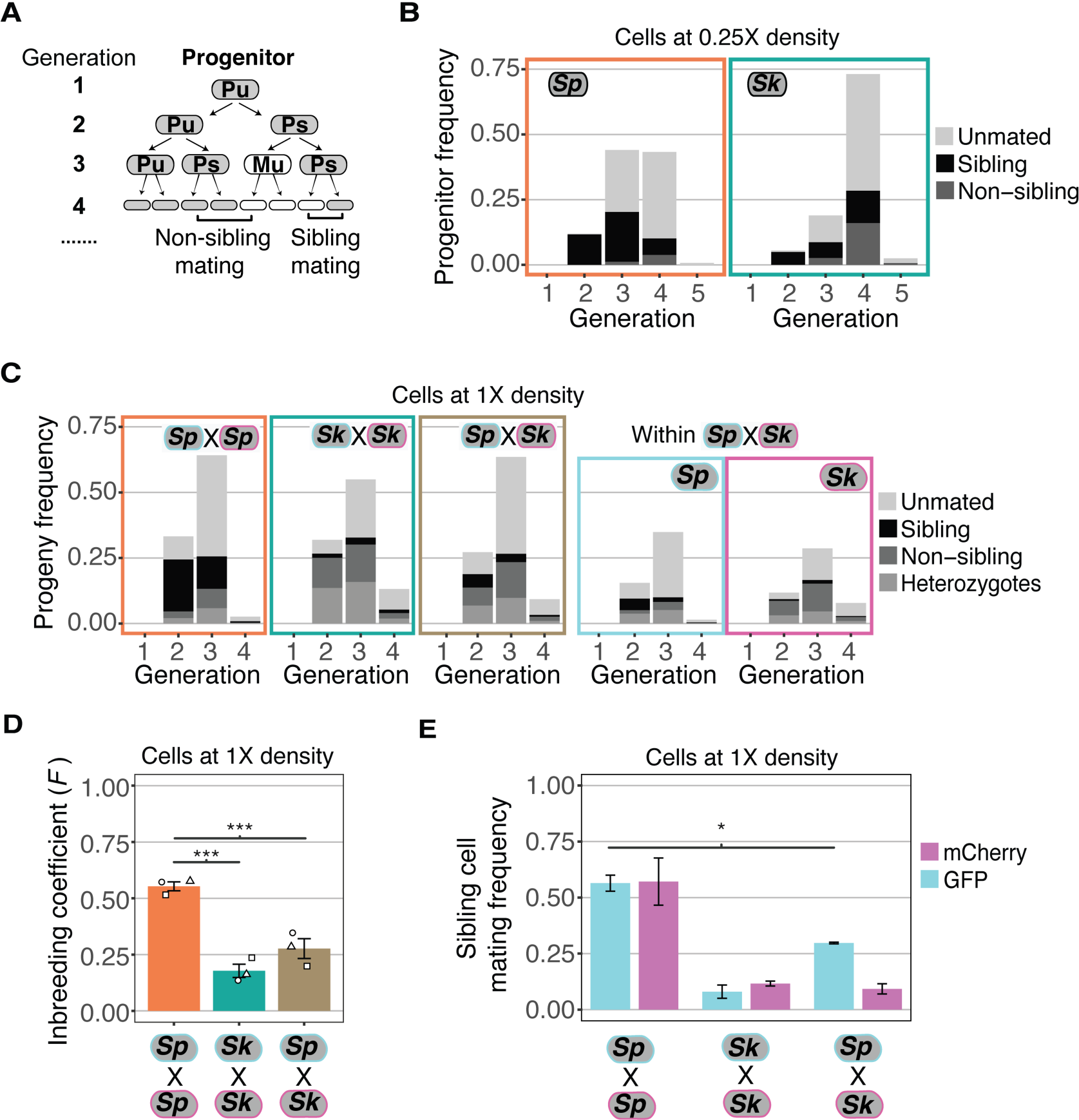
Variation between mating behaviors in *Sp* and *Sk* cells. **A)** Schematic showing cell divisions, switching and mate choice in *Sp*. Mitotic generations are shown on the left. Cells are either *h+* (P) or *h-* (M) and their status is switchable (s) or unswitchable (u). **B)** Division and mating phenotypes of wildtype *Sp* and *Sk* cells plated at low cell density (0.25X) on SPA plates. Single founder cells and their descendants were monitored through time-lapse imaging. Mating patterns were categorized until the end of the generation in which the first mating event occurred. The data presented were pooled from two independent time-lapses in which over 400 founder cells were tracked. **C)** Division and mating phenotypes of fluorescently labeled *Sp* and *Sk* cells plated at standard (1X) density on SPA plates. Individual cells were monitored until the population reached a typical mating efficiency for the specific cross and the mate choice of their progeny and the mitotic generation of those events was classified. The cells labeled with GFP are indicated with a cyan line around the cell whereas the mCherry-labeled cells are outlined in magenta. Pooled data from two independent experiments are presented, with 286 cells scored from each experiment. In the cross of *Sp* (GFP) and *Sk* (mCherry), we present an additional plot (far right) in which the fate of each founder is separated by isolate. **D)** Inbreeding coefficient calculated from still images of cells plated at 1X density on SPA. *** indicates p-value < 0.005, Multiple t-test, Bonferroni corrected. At least three biological replicates per isolate are shown (open shapes). **E)** Breakdown of sibling cell mating by fluorophore in the indicated crosses from C. * indicates p-value=0.04, one-tailed t-test comparing isogenic and mixed *Sp* cells.

Classic work in *Sp* found that only one in a group of four clonally derived cells will have switched mating types. The switched cell will be compatible to mate with one of the other three cells, most typically its sibling (Figure 2A) (Miyata 1981). Under the same switching model, a smaller portion of cells derived from a single division could be of opposite mating types and thus sexually compatible (Figure 2A). In *Sp* cells, we observed mating amongst the clonal descendants of some progenitor cells after a single mitotic division (i.e., at generation 2). By the third generation, we observed mating amongst the descendants of more than half of the progenitor cells. Almost all the observed mating events were between sibling cells (Figure 2A and 2B). These observations are consistent with published work (Klar 1990; Miyata 1981).

*Sk* cells plated at 0.25X density on SPA divided significantly more than *Sp* cells prior to the first mating (Wilcox rank sum test; p< 0.005 Figure 2B). *Sk* progenitor cells most frequently started mating at the fourth mitotic generation (Figure 2B). This phenotype is consistent with less mating-type switching as more generations would be required on average to produce a cell with the opposite mating type (Singh 2002). In addition, many mating events were between non-sibling cells. This phenotype can also be explained indirectly by reduced mating-type switching. Specifically, after cells undergo several divisions, they generate cell clusters in which comparably more non-sibling cells are in close proximity. This clustering could lead to more non-sibling mating than when cells are mating-competent after fewer divisions and the low number of cells are largely arranged linearly.

To better understand the differences between the number of cell divisions prior to mating between *Sp* and *Sk*, we compared the sequence of the mating type locus in the two isolates. Consistent with previous work, we found that the mating-type regions of *Sp* and *Sk* are highly similar (Singh 2002). However, using a previously published mate-pair sequencing dataset we discovered a ∼5 kb insertion of nested *Tf* transposon sequences in the *Sk* mating type region (Eickbush et al. 2019). We confirmed the presence of the insertion using PCR (Supplemental Figure 4A-B). We also found evidence consistent with the same insertion in FY28981, which also mates more randomly than *Sp* (Supplemental Figure 4B, Figure 1D). We did not, however formally test if the insertion affects mating phenotypes. Even if it does have an effect, it is insufficient to explain all the mating type variation we observed as FY29044 mates randomly, yet it lacks the insertion (Figure 1D and Supplemental Figure 4B).

To further explore the hypothesis that decreased mating-type switching efficiency in *Sk* could contribute to the mating differences we observed (Figure 2B), we carried out time-lapse analyses of cells at our standard 1X cell mating density. We reasoned that at this density, any given cell is likely to have a cell of opposite mating type nearby, even if mating-type switching is infrequent. We again used mixed populations of GFP and mCherry-expressing cells to facilitate the scoring of mating patterns (Figure 2C, Supplemental Video 1). We found that the *Sp* cells predominantly mated in the second and third mitotic generations and most mating events were between sibling cells (Figure 2C).

The mating behavior of *Sk* cells changed more dramatically between 0.25x density and the higher 1X density. Whereas *Sk* cells tended to first mate in the fourth generation at 0.25X density, at 1X density *Sk* cells, like *Sp*, generally mated in the second and third mitotic generations (Figure 2C, Supplemental video 2). Additionally, we observed significantly reduced levels of mating between *Sk* sibling cells at 1X density relative to 0.25X (10% and 56%, respectively; Figure 2E and 2B). These phenotypes are consistent with reduced mating-type switching in *Sk*. Specifically, our data suggest that *Sk* cells do not need to undergo more divisions before they are competent to mate. Rather, the additional divisions that occurred at 0.25X density in *Sk* could have been necessary to produce a pair of cells with opposite mating types. At 1X density, additional divisions are not expected to be required as additional non-sibling compatible partners are available.

It is important to note that we were unable to directly measure mating-type switching. Therefore, reduced switching in *Sk* represents a promising model that remains to be tested. Still, our results demonstrate that the mating phenotypes previously measured in *Sp* do not apply to all *S. pombe* isolates. Despite very little genetic diversity, *S. pombe* isolates maintain significant natural variation in key mating phenotypes (Jeffares et al. 2017).

### Ascus variation

While assaying inbreeding cytologically, we noticed that the *Sk* natural isolate displayed tremendous diversity in ascus size and shape (Figure 1C, Supplemental Figure 5A, Supplemental movie 2). This was due to high variability in the size of the mating projections, known as shmoos. *Sk* produced long shmoos only in response to cells of the opposite mating type and not as a response to nitrogen starvation alone (Supplemental Figure 5B). The long *Sk* shmoos motivated us to quantify asci length across all the natural isolates described above. We found that most isolates generated zygotes or asci that were ∼10-15 μm, similar to *Sp*. The majority of *Sk* zygotes and asci also fell within this range, but ∼25% of *Sk* zygotes and asci were longer than 15 μm, with some exceeding 30 μm (Supplemental Figure 5C). We also assayed zygote/ascus length in an additional natural isolate in which we were unable to quantify inbreeding due to a clumping phenotype (FY29033). This isolate also showed populations of long asci, like *Sk* (Supplemental Figure 5C).

Additionally, we occasionally noticed a fused asci phenotype in *Sk* (Supplemental Figure 5D). Time-lapse analyses of mating patterns, described above, revealed these fused asci can result from an occasional disconnect between mitotic cycles and the physical separation of cells (Supplemental Figure 5E). This phenotype is reminiscent of *adg1, adg2, adg3* and *agn1* mutants in *Sp* that have defects in cell fission (Alonso-Nunez et al. 2005; Gould and Simanis 1997; Sipiczki 2007). Although we observed this phenotype in all time lapse experiments using *Sk* cells, the prevalence of this phenotype varied greatly between experiments. We rarely observed this phenotype in *Sp* cells. We did not analyze time-lapse images of the other natural isolates, where this phenotype is most easily observed, so it is unclear if this septation phenotype occurs in other natural isolates.

### Mating phenotypes are affected by available mating partners

We next assayed if the mating preferences of *S. pombe* isolates *Sp* and *Sk* were invariable, or if they could be affected by the available mating partners due to mating incompatibilities (Seike et al. 2019b). To test this idea, we used both still and time-lapse imaging of cells mated at 1X density on SPA. For these experiments, we mixed fluorescently labeled *Sp* and *Sk* cells in equal frequencies.

We observed in experiments employing still images that the overall inbreeding coefficient of the mixed *Sk*/*Sp* population of cells was intermediate between single-isolate crosses and mixed crosses (Figure 2D). In time-lapse experiments, we observed that *Sk* cells maintained low levels of mating between sibling cells in the mixed *Sk*/*Sp* population (9.2% compared to 9.8% in a homogeneous population; Figure 2E). Amongst the *Sp* cells, mating between sibling cells decreased significantly from 56.8% to 29.7% in the mixed mating environment (Figure 2E; t-test, p = 0.04). Together, these results suggest that *Sk* cells can interfere with the ability of *Sp* cells to inbreed. Although sibling cell mating preference changed, we did not observe a significant decrease in the mating efficiency of *Sp* cells in a mixed *Sp/Sk* population relative to a pure *Sp* population (Supplemental Figure 6A). Instead, the mating efficiency in the mixed *Sp/Sk* population was intermediate of those observed in pure *Sp* and *Sk* populations, indicating these isolates do not affect each other’s ability to mate.

We were intrigued by the idea that long shmoos (mating projections) of *Sk* could contribute to its ability to disrupt *Sp* sibling mating. We were unable to address this idea directly. We did, however, find that *Sk*/*Sp* matings produce significantly longer zygotes/asci than either *Sk*/*Sk* or *Sp*/*Sp* matings (Supplemental Figure 6B). This was true even when we compared *Sk*/*Sp* zygote/ascus length to the length of heterozygous *Sk*/*Sk* or heterozygous *Sp*/*Sp* zygotes/asci.

While this result does not prove that long *Sk* shmoos disrupt *Sp* sibling mating, it does show that long shmoos tend to be used in these outcrossing events.

We next extended our analyses by assaying mating efficiency and inbreeding propensity in all pairwise combinations of *Sp*, *Sk*, FY29043, and FY29044 using still images of mated cells. After adjusting for mating efficiencies and parental inbreeding coefficients (see Methods), the phenotypes we observed in these crosses were mostly additive, in that they were intermediate to the pure parental strain phenotypes (Supplemental Figure 7). The two exceptions were in the crosses between *Sk* and the isolates FY20943 and FY20944. *Sk* formed more *Sk*/*Sk* homozygotes than expected in the two crosses (one tailed t-test, p=0.043 and p= 0.038, respectively); suggesting that *Sk* cells may not be fully sexually compatible with FY20943 and FY20944 (Supplemental Figure 7). Overall, our observations indicate that mating phenotypes of a given isolate can be affected by different mating partners. Importantly, however, our results suggest mating incompatibilities are unlikely to have a major role in limiting outcrossing in *S. pombe*.

### Population genetics model of the effect of inbreeding on *wtf* meiotic drivers

We next wanted to test how the observed range of inbreeding values would affect the spread of a *wtf* driver in a population. To do this, we first used population genetic modeling. We used the meiotic drive model presented by J. Crow, but we also introduced an inbreeding coefficient (Hartl and Clark 2007; Crow 1991) (See Methods for a full description of the model). The model considers a population with two possible alleles at the queried locus. We assumed a *wtf* driver would exhibit 98% drive (transmission to 98% of spores) in heterozygotes based on measured values for the *Sk wtf4* driver and other *wtf* meiotic drivers (Bravo Nunez et al. 2020a; Nuckolls et al. 2017). We assumed that homozygotes have a fitness of 1 (e.g., maximal fitness), whereas *wtf* driver heterozygotes have a fitness of 0.51, since meiotic drive destroys nearly half of the spores (Nuckolls et al. 2017). The inbreeding coefficient dictates the frequency of heterozygotes and thus the frequency at which the *wtf* driver can act. We varied the inbreeding coefficient (*F*) from 1 (total inbreeding) to -1 (total outcrossing).

We used the model to calculate the predicted change in the frequency of a *wtf* driver after only one sexual generation (Figure 3A). We also calculated the spread of a *wtf* driver in a population from a 5% starting frequency over many generations of sexual reproduction (Figure 3B). Under complete inbreeding, the frequency of the driver does not increase after sexual reproduction or spread in a population over time (Figures 3A and 3B). No change in driver frequency was expected because no heterozygotes are produced under this condition, so no drive can occur. The *wtf* driver has the greatest advantage if outcrossing is complete. Under all other conditions, including the range of inbreeding coefficients we measured experimentally in *S. pombe* natural isolates, some heterozygotes form and the *wtf* driver increases in frequency over generations of sexual reproduction (Figures 3A and 3B). This model predicts that *wtf* drivers can spread if outcrossing occurs in a population, even if outcrossing is infrequent.

**Figure 3.**
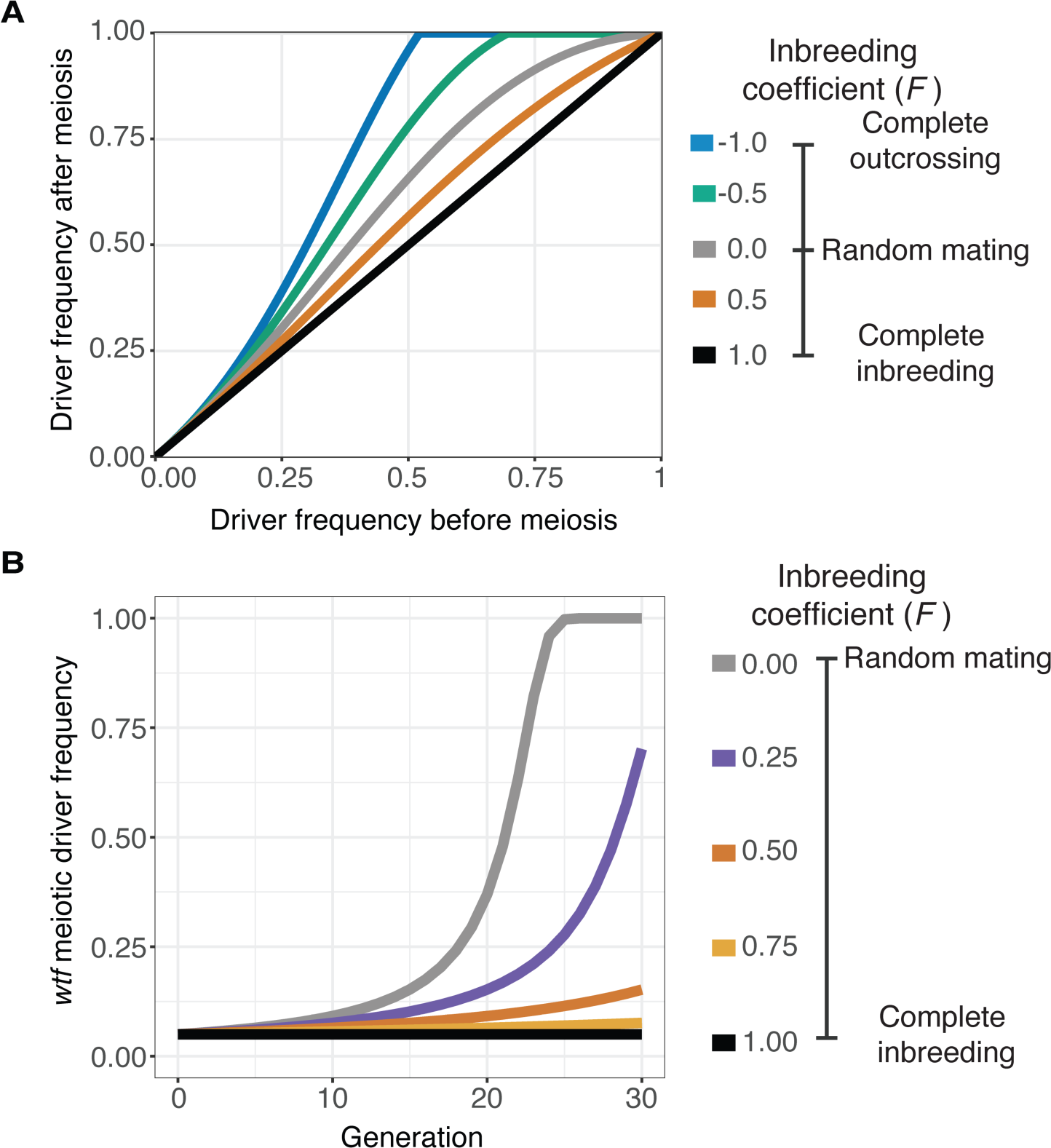
Inbreeding is predicted to slow the spread of a *wtf* driver. We assumed a 0.98 transmission bias favoring the *wtf* driver and a fitness of 0.51 in *wtf+*/*wtf-* heterozygotes. We assumed that homozygote fitness is 1. We simulated the spread of a wtf drive for 30 generations. **A)** Change of *wtf* meiotic driver frequency after meiosis assuming varying levels of inbreeding. **B)** The spread of a *wtf* driver in a population over time assuming varying levels of inbreeding.

### Inbreeding and linked deleterious alleles can suppress the spread of *wtf* drivers

We next wanted to test if our predictions reflect the behavior of *wtf* drive alleles in a laboratory population of *Sp* cells over many generations. To do this, we constructed an experimental evolution system employing the GFP and mCherry fluorescent markers described above to measure changes in allele frequencies in a population over time using cytometry. To mark drive alleles, we linked the fluorescent markers with the *Sk wtf4* driver and integrated the whole construct at the *ura4* locus in *Sp* (Nuckolls et al. 2017). For nondriving alleles, we used GFP or mCherry integrated at the *ura*4 locus without a linked *wtf* gene. We call the non-*wtf* alleles ‘empty vector.’ We started the experimental evolution populations with a defined ratio of GFP and mCherry-expressing cells. We then induced a subset of the population to mate and sporulate followed by collection and culturing of the progeny (spores). From these cells, we remeasured GFP and mCherry frequencies using flow cytometry, and we initiated the next round of mating and meiosis (Figure 4A).

**Figure 4.**
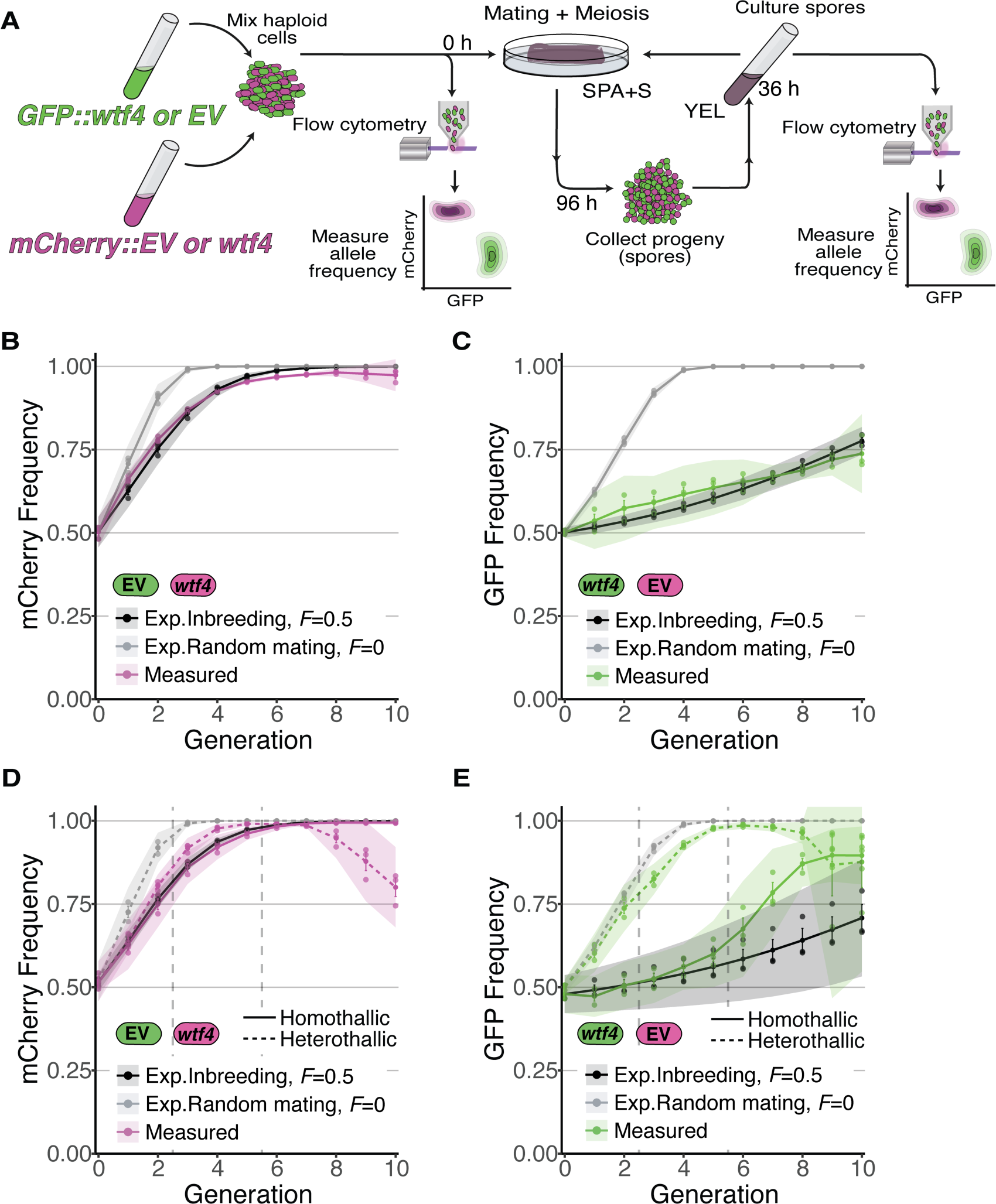
Inbreeding slows the spread of the *wtf4* meiotic driver in homothallic strains. **A)** Experimental strategy to monitor allele frequency through multiple generations of sexual reproduction. GFP (green) and mCherry (magenta) markers are used to follow empty vector (EV) alleles or the *wtf4* meiotic driver. Starting allele frequencies and allele frequencies after each round of sexual reproduction were monitored using cytometry. **B)** Homothallic population with mCherry marking *wtf4* and GFP marking an EV allele. Allele dynamics were predicted using drive and fitness parameters described in the text assuming inbreeding (black lines) and random mating (gray lines). The individual spots represent experimental replicates and the shaded areas around the lines represent 95% confidence intervals. **C)** Same experimental setup as in **B**, but with mCherry marking the EV allele and GFP marking *wtf4*. **D-E)** Repeats of the experiments shown in **B** and **C** with two alterations. First, these experiments tracked heterothallic populations (dotted lines) in addition to homothallic cells (solid lines). Second, the populations were sorted by cytometry at generations 3 and 5 (vertical long-dashed lines) to remove non-fluorescent cells.

Because our experiments rely on comparing the frequency of GFP and mCherry-expressing cells over time, we needed to test the fitness of the markers. We found that both fluorescent markers were lost from all our experimental populations over time (Supplemental Figure 8A-B). This was likely because insertion of the markers disrupted the *ura4* gene and cells that excised the marker reverted the *ura4* mutation and thereby gained a fitness benefit. We therefore only considered fluorescent cells for our analyses and stopped the experiments when more than 95% cells lacked a fluorescent marker. In addition, in one set of experiments we also sorted cells at defined timepoints to remove non-fluorescent cells from our populations (Figure 4D-E and Supplemental Figure 9 C-D, described below).

To assay for potential differences in the fitness costs of GFP and mCherry markers, we carried out our analyses in control populations without drive. One control population lacked the *Sk wtf4* driver while the other had *Sk wtf4* linked to both fluorescent markers. For both types of controls, we analyzed homothallic (inbreeding) and heterothallic (randomly mating) cell populations. We found in most cases that the number of mCherry-expressing cells increased at the expense of GFP-expressing cells over time (Supplemental Figure 9 A-D). The notable exception was in heterothallic populations containing *Sk wtf4* linked to both fluorophore alleles, where we did not observe a different cost of the GFP allele compared to mCherry (Supplemental Figure 9D). The origin of the fitness cost of GFP and why this cost was not observed in the one heterothallic population are both unclear. We do know that the fitness cost of GFP is incurred during sexual reproduction as we observed no differences in vegetative growth between mCherry and GFP- expressing cells (Supplemental Figure 8C-D). We used the first six generations of data from our control crosses (Supplemental Figure 9) to calculate a 11% fitness cost to GFP allele heterozygotes and 22% fitness cost to GFP homozygotes. We then used this cost when predicting the evolutionary dynamics of *wtf4* in the experiments described below (Figure 4B-E).

For our experiments competing *Sk wtf4* with an empty vector allele, we first assayed populations in which the alleles both started at 50% frequency. In homothallic (inbreeding) populations, we observed that *wtf4* alleles spread in the population over several generations of sexual reproduction. The driver spread faster when linked to mCherry than when linked to GFP, presumably due to the aforementioned fitness costs linked to GFP (Figure 4B-C). In both cases, the rate of spread of the allele was very close to our model’s predictions if we assumed an inbreeding coefficient of 0.5 (black lines, Figure 4B-C) and differed considerably from the model’s predictions assuming random mating (gray lines, Figure 4B-C).

We saw similar spread of *wtf4* in homothallic populations in a set of repeat experiments in which we sorted the cell populations twice to remove non fluorescent cells (Figure 4D and 4E). In these experiments, we also assayed heterothallic populations in parallel. As described above, heterothallic cells cannot switch mating type and therefore cannot inbreed. In the heterothallic populations, the *wtf4* driver spread significantly faster than in homothallic cells. In generations 1- 6, the spread of *wtf4* was very similar to that predicted by our model if we assumed random mating. In later generations, our observations did not fit the model well. We suspect extensive loss of fluorescent cells, especially those with mCherry, and the resulting decrease in population size could contribute to this effect (Figure 4D and 4E; Supplemental Figure 8). Overall, our results demonstrate that our population genetics model is good at describing the spread of *wtf4* in our experimental population, particularly in the first few generations. Our results also confirm that incomplete inbreeding slows, but does not stop, the spread of drivers in a population.

We next wanted to further explore the effects of linked deleterious traits on the spread of *wtf* meiotic drivers. We used the population genetic model to calculate the ability of a driver to spread when tightly linked to alleles with fitness costs ranging from 0 to 0.4. We also varied the inbreeding coefficient from 0 (random mating) to 1 (complete inbreeding). We found that in the absence of costs, *wtf* drivers are predicted to spread in a population at all initial frequencies greater than 0 (Figure 5A). As described above, this spread is slowed, but not stopped, by incomplete inbreeding (coefficients less than 1). When the driver is burdened by additional fitness costs, it can still spread in a population. Importantly, the driver must start at a higher initial frequency to spread when linked to deleterious alleles (Figure 5A). For example, when the driver is linked to a locus with a fitness cost of 0.11, like the GFP allele described above, it is expected to spread in a randomly mating population if its initial frequency is 0.125 or higher. In an inbreeding population, the minimal initial frequency required for the driver to spread increases as the degree of inbreeding increases (Figure 5A).

**Figure 5.**
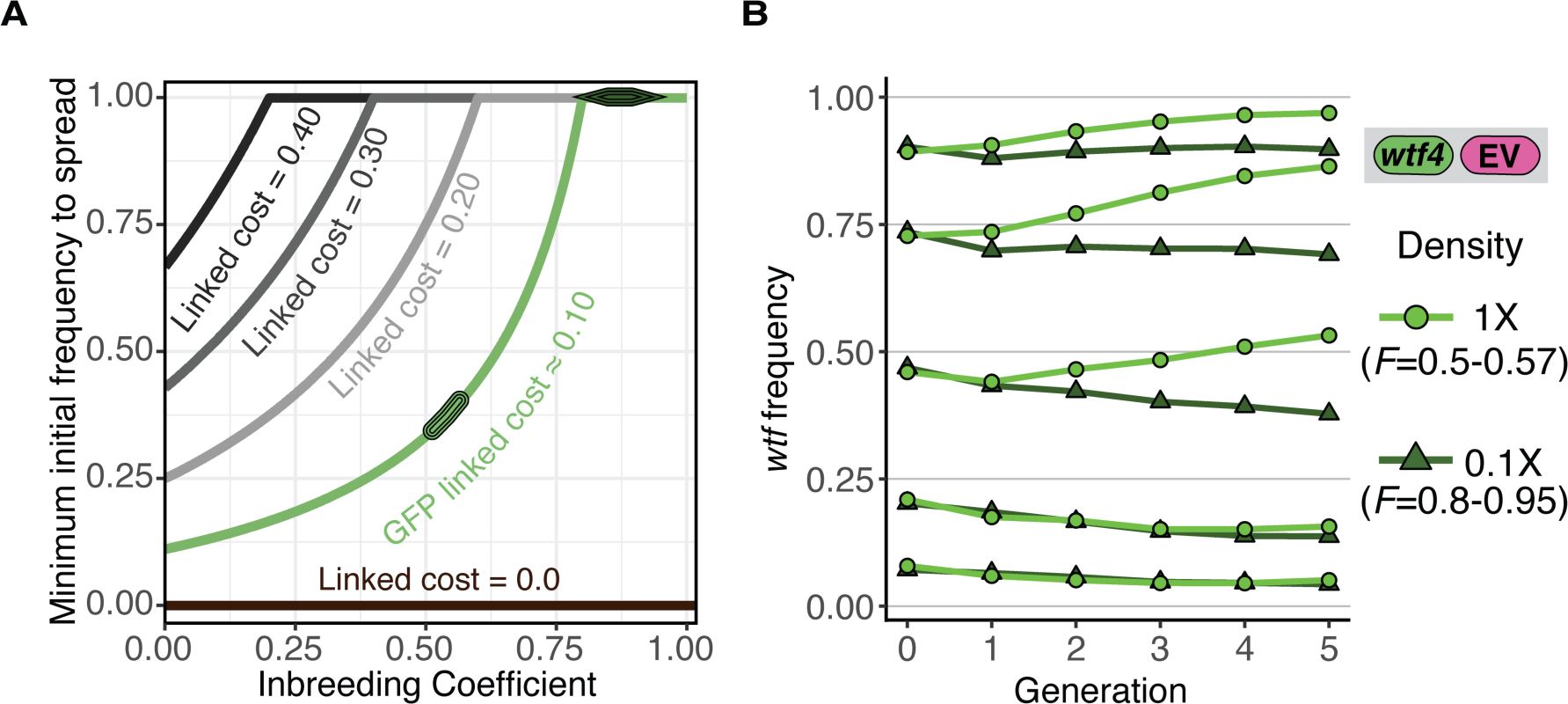
Inbreeding purges *wtf* drivers with linked deleterious alleles. **A)** Modeling the predicted impacts of inbreeding, the fitness of the driving haplotype, and the initial frequency of the driver on the spread of a *wtf* driver in a population. The circle indicates the predicted initial frequency necessary for the GFP::*wtf4*+ allele to spread in a population when mated at standard (1X) density where *F*=0.5. The triangle indicates the predicted initial frequency necessary for the GFP::*wtf4*+ allele to spread in a population when mated at low (0.1X) density where F=0.8-0.95 measured in Supplemental Figure 1C. **B)** Experimental analyses of the impacts of inbreeding and the initial driver frequency on the spread of the GFP::*wtf4*+ allele in a population.

We next tested these predictions experimentally using the *Sk wtf4* allele linked to GFP in homothallic cells. As described above, the GFP allele is linked to an unknown deleterious trait (cost 0.11). We varied the inbreeding of the population by assaying cells mated at 1X and 0.1X density. As reported above, homothallic cells mated at 1X density exhibit an inbreeding coefficient of 0.5 to 0.57, but that is increased to 0.8 to 0.95 by mating the cells at low (0.1X) density (Figure 1D, Supplemental Figure 1B-C). Consistent with the predictions of our model, we observed in the experimental populations that the driver failed to spread when the initial frequency was less than 0.25 (Figure 5B). When the *wtf4:*:*GFP* allele was found in roughly half of the population, it could spread under low levels of inbreeding but decreased in frequency when inbreeding was increased (Figure 5B). Similar, but less dramatic, effects were observed at higher initial frequencies of the *wtf4*::*GFP* allele. Altogether, our experimental analyses are consistent with the predictions of our model and show that both inbreeding and linked deleterious alleles can impede the spread of a *wtf* meiotic driver.

## Discussion

### Natural variation in mating phenotypes in *S. pombe*

Mating phenotypes, particularly the outcrossing rate, are key parameters that affect the evolution of species (Glemin et al. 2019). We sought to explore mating phenotypes in *S. pombe* to better understand the evolution of the *wtf* gene family found in this species. Although genetic variation is limited between *S. pombe* isolates, past studies found variation in mating efficiency and uncovered genetic diversity of the mating-type locus (Jeffares et al. 2015; Nieuwenhuis et al. 2018; Rhind et al. 2011; Singh 2002). In this work, we assayed mating in an array of homothallic *S. pombe* natural isolates under a variety of laboratory conditions. Similar to previous work, we found *Sp* mating efficiency close to 40% and observed variable mating efficiencies for natural isolates (Merlini et al. 2016; Seike et al. 2019a). In addition, we quantified the propensity of natural isolates to inbreed when given the opportunity to outcross. We found this trait was variable between natural isolates and could be affected by cell density or available sexual partners. Finally, we found variation in the size of mating projections and in cell division phenotypes prior to mating.

We did not definitively identify the molecular mechanisms underlying the variation in mating phenotypes we observed. Our data is, however, consistent with a model in which less frequent mating-type switching in the *Sk* isolate contributes to more random mating in *Sk* than in the common lab isolate, *Sp*. Specifically, if switching is less frequent in *Sk*, sibling cells are less likely to be compatible to mate. Incompatibility of sibling cells opens the possibility for mating with other, perhaps non-clonally derived, cells in the population. Singh and Klar were the first to propose that *Sk* switched less frequently than *Sp* when they noticed less of the DNA break that initiates switching (Singh 2002). We discovered a large, nested insertion of transposon sequences in the mating type locus of *Sk,* and we posit that this insertion could contribute to reduced DNA break formation and, potentially, decreased mating-type switching. The long shmoos we observed in *Sk* may also contribute to more random mating in this isolate, as the long shmoo may increase the available number of partners within range.

Additional previously described natural variation that we did not functionally explore may also contribute to differences in inbreeding propensity in *S. pombe*. For example, heterothallic natural isolates are predicted to exclusively outcross to isolates with the opposite mating type (Jeffares et al. 2015; Nieuwenhuis et al. 2018). In addition, homothallic isolates with atypical mating type loci with extra copies of the *mat* cassettes could grow into populations that are biased towards one mating type (Nieuwenhuis and Immler 2016). Indeed, we analysed the presumably expressed mating type locus (*mat1*) in several isolates for which we had nanopore sequencing data and found an approximate 3:1 excess of the *h+* allele in FY29033 (Supplemental figure 4C). The excess of one mating type is predicted to facilitate outcrossing.

It is important to note that our study does not address the actual frequency of outcrossing in *S. pombe* populations in the wild. Very little is known about the ecology of fission yeast, including how frequently genetically distinct isolates are found in close enough proximity to mate (e.g., closer than ∼40 microns apart) (Jeffares 2018). Outcrossing rates have been estimated using genomic data, but those estimates generally assume both that heterozygous recombination rates will match those observed in pure *Sp* and that allele transmission is Mendelian (Farlow et al. 2015). Although these genomic estimates are reasonable, neither of these assumptions is consistent with empirical analyses (Bravo Nunez et al. 2020b; Hu et al. 2017; Zanders et al. 2014). These assumptions have, therefore, likely led to an underestimation of the true outcrossing rate.

### The effect of mating-type phenotypes on the spread of *wtf* meiotic drivers

To understand the evolution of the *wtf* drive genes, it is not necessarily essential to understand how frequently significantly diverged natural isolates mate. Instead, it is important to understand how often a driver is found in a heterozygous state. This is likely a significantly higher frequency than the frequency of mating between diverged isolates because of the rapid evolution of the *wtf* gene family. Even though genetic diversity within *S. pombe* is low (<1% average DNA sequence divergence in non-repetitive regions), the *wtf* genes present in different isolates tend to be largely distinct (Eickbush et al. 2019; Hu et al. 2017; Jeffares et al. 2015; Rhind et al. 2011). The number of *wtf* genes per isolate varies from 25-38 *wtf* genes (including pseudogenes), and even genes found at the same locus can be dramatically different (e.g., <61% coding sequence identity between alleles of *wtf24*) (Eickbush et al. 2019). The rapid evolution of *wtf* genes is driven largely by non-allelic gene conversion within the family and expansion or contraction of repetitive sequences within the coding sequences of the genes (Eickbush et al. 2019).

Importantly, *wtf* genes generally provide no protection against *wtf* drivers with distinct sequences (Bravo Nunez et al. 2018; Bravo Nunez et al. 2020a; Hu et al. 2017). As a result, even small sequence changes in *wtf* drivers can cause the birth and death of drivers. When a cell bearing a novel *wtf* driver mutation mates with a cell without the mutation (i.e., not with its identical sibling cell), the driver is heterozygous, and thus, drive can occur.

Previous work assayed the strength of drive and the associated fitness costs of *wtf* drivers (Bravo Nunez et al. 2018; Bravo Nunez et al. 2020a; Hu et al. 2017; Nuckolls et al. 2017). Those data, along with the inbreeding coefficients measured in this study, allowed us to mathematically model the spread of a *wtf* meiotic driver in a *S. pombe* population. Our modeling showed that the incomplete inbreeding we observed in *S. pombe* could slow the spread of a *wtf* driver. Importantly, the incomplete inbreeding observed in *S. pombe* does not halt the spread except in cases where the driver is found in low frequencies and linked to a deleterious allele. Given the tractability of *S. pombe*, we were also able to test the predictions of the model experimentally. Overall, our experimental results were quite similar to the model’s predictions discussed above. This suggests that our model encompasses all critical parameters. In addition, our experiments show how the *wtf* drivers can persist and spread in *S. pombe*, even if outcrossing is infrequent. The variation of mating phenotypes also indicates that the rate of spread of a *wtf* driver is expected to vary between different populations of *S. pombe*.

Overall, our results are consistent with previous empirical and modeling studies of meiotic driver dynamics in populations. For example, like our fortuitously deleterious GFP allele, meiotic drivers are often linked to deleterious alleles that can hitchhike with the driver (Atlan et al. 2004; Dyer et al. 2007; Finnegan et al. 2019; Fishman and Saunders 2008; Higgins et al. 2018; Lyon 2003; Olds-Clarke 1997; Schimenti et al. 2005; Wilkinson and Fry 2001; Wu 1983). The added costs reduce the spread of drivers, which can lead a population to harbor a driver at stable intermediate frequency (Dyer and Hall 2019; Finnegan et al. 2019; Fishman and Kelly 2015; Hall and Dawe 2018; Manser et al. 2011).

### *S. pombe* as a tool to experimentally model complex drive dynamics

To conclude, we would like to highlight the potential usefulness of the *S. pombe* experimental evolution approach developed for this study. With this system, we were able to observe the effects of altering allele frequencies, inbreeding rate, and fitness of a driving haplotype. In the future, this system could be used to experimentally explore additional questions about drive systems. For example, one could experimentally model meiotic drivers that bias sex ratios by linking the driver to the mating type locus in a heterothallic population. In addition, one could explore the evolution of complex multi-locus drive systems employing combinations of multiple *wtf* meiotic drivers or drivers and suppressors. This tool could lead to novel insights about natural drivers, but it may also be particularly useful for exploring potential evolutionary trajectories of artificial gene drive systems (Burt and Crisanti 2018; Price et al. 2020; Wedell et al. 2019).

## Materials and methods

### Generation of *ura4*-integrating vectors and fluorescent strains

We introduced the fluorescent genetic markers into the genome using plasmids that integrated at the *ura4* locus. To generate the integrating plasmids, we first ordered gBlocks from IDT (Coralville, IA) that contained mCherry or GFP under the control of a *TEF* promoter and *ADH1* terminator (Hailey et al. 2002; Sheff and Thorn 2004). We digested the gBlocks with SpeI and ligated the GFP cassette into the SpeI site of pSZB331 and the mCherry cassette into the SpeI site of pSEZB332 (alternate clone of pSZB331; (Bravo Nunez et al. 2020a; Bravo Nunez et al. 2020b)) to generate pSZB437 and pSZB882, respectively. We then linearized the plasmids with KpnI and transformed them into *S. pombe* using the standard lithium acetate protocol (Schiestl and Gietz 1989). We independently transformed the isolates GP50 (*S. pombe*), *S. kambucha*, FY28974, FY28981, FY29022, FY29033, FY29043, and FY29044. We were unsuccessful in transforming FY28969, FY29048, and FY29068. FY29033 was not included in the inbreeding analyses due to its proclivity to clump. The homothallic and heterothallic strains carrying mCherry or GFP were transformed using the same method.

To add *Sk wtf4* to the *Sp* genome, we again used a *ura4*-integrating plasmid. To generate this plasmid, we amplified *Sk wtf4* from SZY13 using the oligos 688 and 686. We digested the amplicon with SacI and ligated into the SacI site of pSZB332 to generate pSZB716 (Bravo Nunez et al. 2020a; Bravo Nunez et al. 2020b). We then separately introduced the GFP and mCherry gBlocks into the SpeI site of pSZB716 to generate pSZB904 and pSZB909, respectively. We introduced the resulting plasmids into yeast as described above.

### Crosses

We performed crosses using standard approaches (Smith 2009). We cultured each haploid parent to saturation in 3 ml YEL (0.5% yeast extract, 3% dextrose, and 250 mg/L adenine, histidine, leucine, lysine, and uracil) for 24 hours at 32°C. We then mixed an equal volume of each parent (700 μL each for individual homothallic strain, 350 μL for heterothallic parents), pelleted and resuspended in an equal volume of ddH_2_O (1.4 mL total), then plated 200 μL on SPA (1% glucose, 7.3 mM KH_2_PO_4_, vitamins and agar) for microscopy experiments or SPA+S (SPA + 45 mg/L adenine, histidine, leucine, lysine and uracil) for genetics experiments. We incubated the plates at 25°C for 1-4 days, depending on the experiment (see figure legends for exact timing). When we genotyped spore progeny, we scraped cells off of the plates and isolated spores after treatment with B-Glucuronidase (Sigma) and ethanol as described in (Smith 2009).

### Iodine staining

We grew haploid isolates to saturation in 3 ml YEL overnight at 32°C. We washed the cells once with ddH_2_O then resuspended them in an equal volume ddH_2_O. We then spotted 10 μL of each strain onto an SPA+S plate, which we then incubated at 25°C for 4 days prior to staining with iodine (VWR iodine crystals) vapor (Forsburg and Rhind 2006).

### Mating-type locus assembly and PCR

We used mate-pair Illumina sequencing reads to assemble the mating-type locus of *S. kambucha* with previously published data (Eickbush et al. 2019). We assembled the mating-type locus using Geneious Prime software, (https://www.geneious.com; last accessed March 18, 2019) using an analogous approach to that described to assemble *wtf* loci (Eickbush et al. 2019).

### DNA extraction for Nanopore sequencing

To extract DNA for Nanopore sequencing we used a modified version of a previously developed protocol (Jain et al. 2018). We pelleted 50 ml of a saturated culture and proceeded as described, with the addition of 0.5 mg/ml zymolyase to the TLB buffer immediately prior to use.

### Nanopore sequencing and assembly

We used a MinION instrument and R9 MinION flow cells for sequencing. For library preparation, we used the standard ligation sequencing prep (SQK-LSK109), including end repair using the NEB End Prep Enzyme, FFPE prep using the NEB FFPE DNA repair mix, and ligation using NEB Quick Ligase. We did not barcode samples and thus used each flow cell for a single genome. We used guppy version 2.1.3 for base calling. We removed sequencing adapters from the reads using porechop version 0.2.2 and then filtered the reads using filtlong v0.2.0 to keep the 100x longest reads. We then error corrected those reads, trimmed the reads and de novo assembled them using canu v 1.8 and the ovl overlapper with a predicted genome size of 13 mb and a corrected error rate of 0.12 (Koren et al. 2017). Base called reads are available as fastq files at the SRA under project accession number PRJNA732453.

In order to count allele frequency within the active *mat* locus, we mapped raw reads back to the corresponding de novo assembly using graphmap v0.5.2 and processed using samtools v1.12 (Li et al. 2009; Sović et al. 2016). We then visually observed the reference-based assemblies using IGV v2.3.97 to count the number of *h+* and *h-* alleles present at the active mating type locus with anchors to unique sequence outside the *mat* locus (Robinson et al. 2011).

### Measuring inbreeding coefficients by microscopy

We mixed haploid parents (a GFP and an mCherry-expressing strain) in equal proportions on SPA as described above. We then left the plate to dry for 30 minutes and then took a punch of agar from the plate using a 1271E Arch Punch (General Tools, Amazon). We then inverted the punch of agar into a 35 mm glass bottomed dish (No. 1.5 MatTek Corporation). We used this sample to count the initial frequency of the two parental types. We then imaged a second punch of agar taken from the same SPA plate after 24 hours incubation at 25°C for homothallic cells and 48 hours for heterothallic cells.

To image the cells, we used an AXIO Observer.Z1 (Zeiss) wide-field microscope with a 40x C-Apochromat (1.2 NA) water-immersion objective. We excited mCherry using a 530–585 nm bandpass filter which was reflected off an FT 600 dichroic filter into the objective and collected emission using a long-pass 615 nm filter. To excite GFP, we used a 440-470 nm bandpass filter, reflected the beam off an FT 495 nm dichroic filter into the objective and collected emission using a 525-550 nm bandpass filter. We collected emission onto a Hamamatsu ORCA Flash 4.0 using µManager software. We imaged at least three different fields for each sample.

We used cell shape to identify mated cells (zygotes and asci) and used fluorescence to identify the genotype of each haploid parent. To measure both fluorescence and the length of asci, we used Fiji (https://imagej.net/Fiji) software to hand draw 5 pixel-width lines through the length of each zygote or ascus. After subtracting background using a rolling ball background subtraction with width 50 pixels, we then measured the average intensity for the GFP and mCherry channels. When measuring the log10 ratio of GFP over mCherry, the mCherry homozygotes have the lowest ratio, homozygotes for GFP the highest ratio, and heterozygotes intermediate.

To calculate the inbreeding coefficient, we used the formula F=1- (observed heterozygotes/expected heterozygotes). We used Hardy-Weinberg expectations to calculate the expected frequency of heterozygotes (2*p*(1-*p*)) for each sample, where ‘*p*’ is the fraction of mCherry+ cells and (1-*p*) is the fraction of GFP+ cells measured prior to mating (Hartl and Clark 2007).

### Visualizing mating and meiosis using time-lapse microscopy

For time-lapse imaging of cells mated at 1X density (Figure 2C), we prepared cells using the agar punch method described above. For cells at 0.25X density (Figure 2B), we used the same approach, except we cultured cells in 3 mL EMM (14.7 mM C_8_H_5_KO_4_, 15.5 mM Na_2_HPO_4_, 93.5 mM NH_4_Cl, 2% w/v glucose, salts, vitamins, minerals) then washed three times with PM-N (8mM NA_2_HPO_4_, 1% glucose, EMM2 salts, vitamins and minerals) before plating cells to SPA. While imaging the cells, we added a moistened kimwipe to the MatTek dish to maintain humidity. We sealed the dish lids on with high-vacuum grease (Corning). We imaged cells using either a Ti Eclipse (Nikon) coupled to a CSU W1 Spinning Disk (Yokagawa), or a Ti2 (Nikon) widefield using the 60x oil immersion objective (NA 1.45), acquiring images every ten minutes for 24-48 hours, using a 5x5 grid pattern with 10% overlap between fields. The Ti Eclipse was used for one replicate each of the 1x crosses and the Ti2 was used for all remaining experiments. We used an Okolab stage top incubator to maintain the temperature at 25°C. For the Ti2 (widefield) data we excited GFP through a 470/24 nm excitation filter and collected through an ET515/30m emission filter. For mCherry on this system, we excited through a 550/15 nm excitation filter and collected through an ET595/40m emission filter. For the Ti Eclipse (confocal) data, we excited GFP with a 488 nm laser and collected its emission through an ET525/36m emission filter. For mCherry on this system, we excited with a 561 nm laser and collected through an ET605/70m emission filter.

To monitor mating in 1X crosses (Figure 2C and 2E), we recorded the number of divisions and mating choice of the progeny of 286 cells until an expected mating efficiency for the population being filmed was attained. The expected mating efficiency was calculated from still images of the same crosses. We recorded two videos of each cross.

To monitor the number of divisions required before mating could occur in 0.25X cultures (Figure 2B), around 200 individual cells were monitored through the duration of the generation in which the first mating event occurred. If cells failed to mate, they were monitored throughout the duration of the movie. If a cell or its mitotic offspring interacted with a neighboring cell cluster, it was not included in the analysis. We recorded two videos for each isolate.

### Calculation of mating efficiency

We calculated mating efficiency from microscopic images using the following formula:

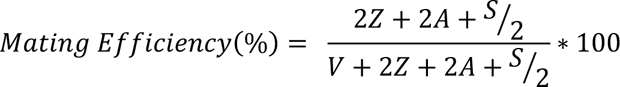

Where *Z* represents the number of zygotes, *A* represents the number of asci, *S* represents the number of free spores and *V* represents the number of vegetative cells (Seike and Niki 2017).

### Measuring inbreeding coefficients using genetics

We used a high-throughput system to genotype the progeny from each cross. First, we crossed two parental populations to generate spore progeny as described above. In addition to placing the mixed haploid cells on SPA+S, we also diluted a subset of the mix and plated it onto YEA+S. We genotyped the colonies that grew on that YEA+S plate to measure the starting frequency of each parental strain in the cross. We plated the spores produced by the cross on YEA+S and grew them at 32°C for 4 days. We picked the colonies using a Qpix 420 Colony Picking System and cultured them in YEL for 24 hours at 30°C in 96 well round-bottom plates (Axygen). We then used a Singer RoTor robot to spot the cultures to YNP dropout and YEA+S drug plates and incubated them at 32°C for three days. We then imaged the plates using an S&P robotics SPImager with a Canon EOS Rebel T3i camera. We analyzed each picture using the *subtract background* feature in Fiji and assigned regions of interest (ROIs) to the 384 spots where cells were pinned. We then measured the average intensity of each spot and classified cells as grown or not by a heuristic threshold. We genotyped some cross progeny manually using standard techniques due to robot unavailability, with indistinguishable results.

We then inferred the frequency of outcrossing based on the frequency of recombinant progeny using a combination of either two or three unlinked genetic markers. If mating was random, we expect the progeny to reflect Hardy-Weinberg expectations (*p*^2^ + 2*p*(1 − *p*) + (1 − *p*)^2^ = 1), where *p*^2^ + (1 − *p*)^2^ reflect the expected frequency of homozygotes and 2*p*(*1-p*) reflects the expected frequency of outcrossing. If the parental strains inbreed to make homozygotes, they can only produce offspring with the parental genotypes. If the strains outcross to generate heterozygotes, they will make the parental genotypes and recombinant genotypes all in equal frequencies (2^n^ total genotypes where n is the number of segregating markers). For our crosses with three markers, we therefore expected the true ‘observed’ frequency of progeny produced by outcrossing to be equal to the number of observed recombinants divided by 6/8. For our crosses with two unlinked markers, we divided by 2/4. We then calculated the inbreeding coefficient using the formula,

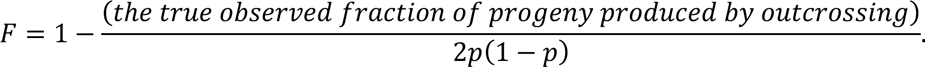

### Zygote frequency expectation under inbreeding and different mating efficiencies

We calculated the expected zygote frequencies when isolates were outcrossed with different isolates on SPA (see Measuring inbreeding coefficients by microscopy) using an additive model that incorporated mating efficiencies and inbreeding coefficients measured from the isogenic crosses. The model assumed that each strain contributes equally to inbreeding, and that they do not change their own mating in response to the mating partner.

We calculated the expected frequency of homozygotes for parental strain 1 as:

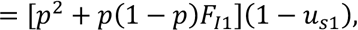

where the inbreeding coefficient is *F*_*I*1_ and the mating efficiency is (1 − *U*_*s*1_) for parental strain 1 (*s*1), considering its initial frequency *p*. The expected heterozygote frequency is:

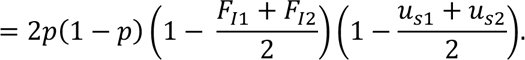

The expected fraction of homozygotes for parental strain 2 is:

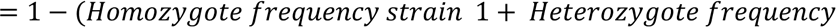

### Calculating expected allele frequencies after sexual reproduction

To model the expected changes in allele frequencies in a randomly mating population over time, we used the equations described by Crow (Crow 1991). For nonrandomly mating populations, we included the ‘*F*’ inbreeding coefficient in the equations, similar to (Sun and Tian 2017). We assumed that *wtf4* had a drive strength (‘*k*’) of 0.98 based on experimental observations (Nuckolls et al. 2017). We assumed the control allele exhibited Mendelian transmission (*k*)=0.5.

Simulations for the spread of a driver in Figure 3 only considered drive and inbreeding. To simulate drive in fluorescent populations, the starting frequencies of each allele (i.e. ‘*p*’) were determined empirically for each experiment using either traditional genetic approaches or cytometry. For relative fitness values *W*_11_, *W*_12_, *W*_22_, we assigned mCherry/mCherry homozygotes a fitness of *W*_11_ = 1, regardless of whether they were EV/EV or *wtf4*/*wtf4* homozygotes. In all but one cross (see below), we observed a fitness cost linked to the GFP alleles relative to mCherry alleles during sexual reproduction, regardless of *wtf4*. We therefore used our data (see below) to calculate the 0.11 as the fitness cost of the GFP-linked variant. Because of that, we assigned a fitness value of 0.78 to GFP homozygotes, *W*_!!_. We assumed the fitness cost linked to GFP was codominant and thus assigned a fitness value of 0.89 for GFP/mCherry homozygous for *wtf4* or Empty vector, *W*_#!_. For GFP/mCherry heterozygotes that were also heterozygous for *wtf4*, we assigned a fitness of 0.46, *W*_#!_. This accounts for the cost of spores killed by drive (0.5/0.98=0.51) and the cost of the GFP-linked variant (0.51*0.89=0.46) The calculation for allele frequency for a *wtf* meiotic driver in consecutive sexual cycles from haploid populations is,

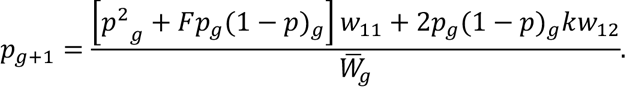

Where the mean fitness of the population at each generation (*g*),

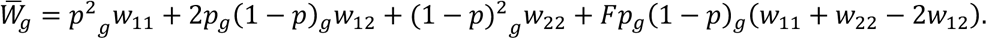

To determine the fitness cost of the GFP linked variant that was present in most of our experiments, we used the L-BFGS-B algorithm to find a fitness that maximized the likelihood that our observed allele frequencies varied only due to the cost of GFP in the first six generations of the control experiments shown in Supplemental Figure 9 (Byrd et al. 1995). To do this, we used the *mle* function from the R *stats4* package (Team 2019). We assumed the fitness cost alters the relative fitness of both homozygotes and heterozygotes. The fitting was done using only the initial six generations due to a rapid loss of fluorescent cells from seven to ten generations.

To calculate the minimum initial frequency of driver linked to alleles with varying additional fitness costs, we assumed codominance for the additional alleles (*c*). Then the relative fitness of heterozygotes for the allele is *W*_12_ = 1 − *c* and the relative fitness of homozygotes for the allele is *W*_22_ = 1 − 2*c*. For simplicity we also assumed complete transmission bias, *k* = 1. The relative fitness of the alternative allele was assumed to be *W*_22_ = 1. A *wtf* meiotic driver linked to deleterious allele spread under the condition that

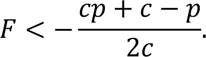

Where *p* is the initial frequency of a *wtf* driver linked to allele *c* and *F* was the inbreeding coefficient.

### Measuring allele frequencies for experimental evolution analyses

We performed the crosses and collected spores as described above. We then started the next generation by culturing 60 µL of spores from each cross in each of three different wells with 600 µL fresh YEL media in 96 deep-well-round-bottom plates (Axygen) and cultured for 24 hours at 1200 rpm at 32°C. We then transferred 60 µl from each culture to a new plate with 600 µL YEL and cultured for 12-14 hours at 1200 rpm at 32°C. We then pooled the culture replicates in an Eppendorf tube, spun down, and resuspended them in an equal volume of ddH20. We then took 100 µL of this sample to assay via cytometry (described below). We also plated 200 µL of each sample on SPA+S plates and incubated at 25°C for 5 days to allow the cells to mate and sporulate.

To detect and quantify fluorescent cells via flow cytometry, we used the ZE5 Cell Analyzer (Bio-Rad). We spun down 100 µL of each culture, washed the cell pellet with water, spun down again and resuspended the cells in 200 µL of 1X PBS (phosphate buffered saline) with 1.5 μl 4′,6- diamidino-2-phenylindole (DAPI, Sigma-Aldrich, 100 ng/ml). DAPI stains dead cells, so we considered DAPI-negative cells as live cells. To image DAPI, we used 355 nm laser excitation and a 447/60 nm detector. To image GFP, we excited using a 488 nm laser and detected emission with a 525/35 nm filter. To image mCherry, we excited using a 561 nm laser and detected emission with a 615/24 nm filter. We used 405 nm and 488 nm (FSC 405 and FSC 488) lasers for forward scatter and 488 nm laser for side scatter (SSC 488).

To quantify the frequency of GFP or mCherry positive cells, we analyzed the Flow Cytometry Standard files in R/Bioconductor using the packages FlowTrans and FlowClust (Lo et al. 2009). We first separated the cells that had round and uniform shape and similar granularity. This step allowed us to detect a more uniform population of single cells. We then discarded DAPI positive cells. To determine the GFP positive and mCherry positive cells, we used limits for each channel. The limits varied throughout the experiment due to reconfiguration in the flow cytometer. For every measurement, we corrected with standard samples of cells that only expressed either GFP or mCherry. Fluorescent GFP positive and mCherry positive cells always showed non-overlapping cell populations. We quantified cells that were not classified as non-fluorescent cells.

### Fluorescent marker loss in experimental evolution

In the experimental evolution experiments (Figure 4, Figure 5B, and Supplemental Figure 9), a large proportion of cells lost their fluorescent marker over time (Supplemental Figure 8 A-B). We assume this is because we introduced the markers using an integrating plasmid that enters the genome following a single crossover event. Because of this, a single crossover can then pop the marker and the associated vector out of the genome. We repeatedly observed that the loss occurred faster with the mCherry markers. Consistent with this model, cells that lost fluorescence generally also lost the associated drug resistance marker present on the integrating vector.

We only considered fluorescent cells in our analyses. In the experiments (Figure 4A-B, Supplemental Figure 4A-B, and Figure 5B) we extended the evolution up to ten generations. In other experiments (Figure 4C-D and Supplemental Figure 4C-D), we removed non-fluorescent cells by cell sorting after generations two and five. To sort cells, we first collected and cultured spores as described above with the first culture done for 12 hours. We then transferred 60 µL of germinated cells into 600 µL fresh YEL media. We used three cultures for each experimental line and pooled all in equal amounts to have 1.4 ml of each line. We spun each sample down and resuspended the cells in 5 ml ddH_2_0 prior to sorting. We removed non-fluorescent cells and retained GFP positive (488 nm laser for excitation and a filter 507 nm) and mCherry positive (561 nm laser for excitation and a 582 nm filter) cells using the laser the BD FACSMelody cell sorter software. We collected 1.2 million cells for each line into 1X PBS. We then spun down the cells and resuspended the pellets in 200 µL YEL. We then took 60 µL from each sample and diluted the cells into 600 µL YEL and continued with the time course as described above. This restored fluorescently labelled cells populations as expected for homothallic and heterothallic lines (Supplemental Figure 8B). Cell sorting did not affect our results as we observed the same patterns in replicate experiments in which we did not remove the non-fluorescent cells (Figure 4).

## Acknowledgements

We thank members of the Zanders lab, María Bravo Núñez and Ibrahim M. Sabbarini for their helpful comments on the paper. We are grateful to Gerry Smith for sharing various strains and thank Alexandra Cockrell for technical support. Original data underlying this manuscript can be accessed from the Stowers Original Data Repository at http://www.stowers.org/research/publications/libpbxxxx. This work was performed to fulfill, in part, requirements for JLH’s thesis research in the Graduate School of the Stowers Institute for Medical Research. This work was supported by The Stowers Institute for Medical Research (SEZ); the Searle Scholars Award (SEZ); and the National Institutes of Health (NIH) DP2GM132936 (SEZ). The funders had no role in study design, data collection and analysis, or manuscript preparation. The content is solely the responsibility of the authors and does not necessarily represent the official views of the funders.

## Data availability

Supplemental file 1. *S. pombe* natural isolates.

List of *S. pombe* natural isolates reported by (Tusso et al. 2019). Those selected to measure inbreeding coefficients using microscopy are highlighted and observations for these from other studies are reported (Jeffares et al. 2015; Nieuwenhuis et al. 2018).

Supplemental file 2. Raw data of allele transmission values reported in Supplemental figure 1.

Crosses shown in Supplemental figure 1 were genotyped before and after meiosis. The absolute and relative frequencies of each genotype were quantified. Segregation for each genetic marker is reported. From each cross, the list of recombinant genotypes are described. In tables 1-3 on spreadsheet 1 (columns B-K) we show the genotypes, number of colonies counted and initial frequencies of the two parents used in each cross represented in Supplemental figure 1B. In columns L, O and R we show the expected frequency of Parent1, Parent 2 and recombinants, respectively, predicted using Hardy-Weinberg. Columns M-Y show the results obtained after meiosis. In columns M-N, P-Q and S-T we report the observed frequency and total number of colonies counted for Parent 1, Parent2, and recombinants, respectively. In columns U-W (U-X for table 3) we report the transmission frequency of each individual marker in the recombinants. The inbreeding coefficient for each cross is reported in Column Y. In tables 4,5 and 6 we show the numbers of each genotype obtained after meiosis for each replicate experiment. Parental genotypes are shown in white. Discordant genotypes are highlighted in blue at the bottom of each table. We also specify which experiments were performed manually and which were performed using robotics. Spreadsheet 2 shows similar data for the multiple density experiments shown in Supplemental figure 1C-D. Column Q also shows the G of fit G-test p-value.

Supplemental file 3. Yeast strains used in this study.

Strains used and created in this study are listed.

Supplemental file 4. Plasmids used in this study.

Plasmids used in this study.

Supplemental file 5. Oligo table.

Oligos used in this study.

Supplemental video 1. Homothallic *Sp* mating video.

Homothallic *Sp* cells plated mating inducing media (SPA) were recorded for 24 hours. Cells labelled with constitutively -expressing fluorophores mCherry (magenta) or (GFP) were mixed in equal proportion 1X standard cell density.

Supplemental video 2. Homothallic *Sk* mating video.

Homothallic *Sk* cells plated mating inducing media (SPA) were recorded for 24 hours. Cells labelled with constitutively -expressing fluorophores mCherry (magenta) or (GFP) were mixed in equal proportion at 1X standard cell density.

---

**Supplemental figure 1.**
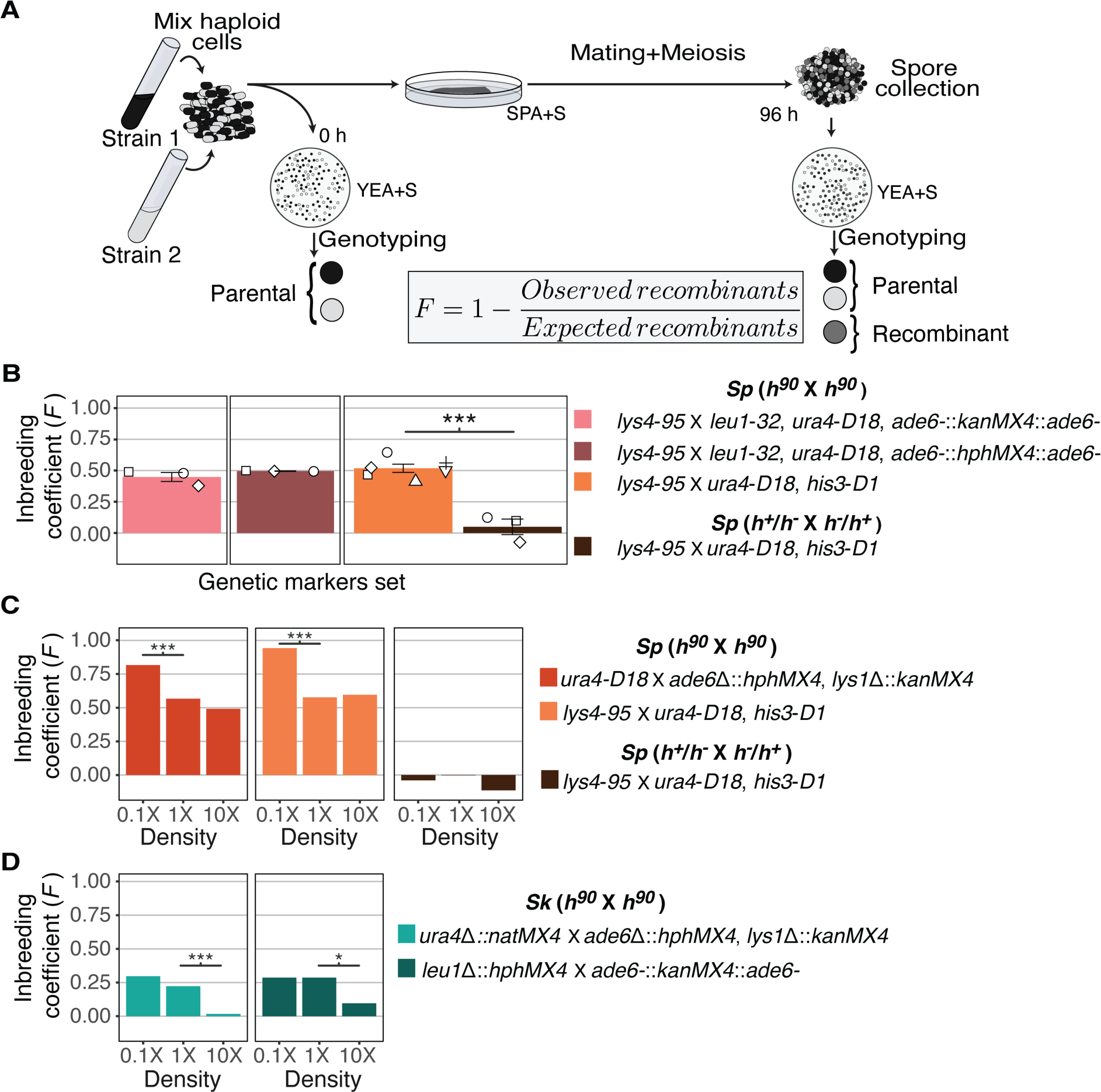
Inbreeding coefficients can be affected by cell density. **A)** Experimental strategy to infer inbreeding coefficient by genotyping the segregation of unlinked genetic markers. The composition of the starting population was measured by genotyping cells placed on rich medium (YEA+S). Cells were also plated on SPA+S, where they mate and undergo meiosis. After meiosis, spores were genotyped and we compared the expected number of recombinant progeny to the number observed to calculate the inbreeding coefficient (*F*). **B)** Inbreeding coefficients calculated as described in **A** using the indicated genetic markers at standard 1X mating density. Each color indicates a different cross. *** indicates p-value<0.005, One tailed t-test on at least three biological replicates (open shapes). **C**) Inbreeding coefficients calculated as described in **A** for *Sp* crosses plated at low (0.1X), standard (1X), and high (10X) density. *** indicates p-value <0.005, G-test. Colonies were randomly sampled for each cross–320 for left panel and 144 colonies for the second and third cross sets. **D)** Inbreeding coefficients calculated as described in **A** for *Sk* crosses plated at low (0.1X), standard (1X), and high (10X) density. *** indicates p-value <0.005, *indicates p-value < 0.05 G-test. 325 and 340 colonies were sampled from the crosses in the left and right panels, respectively.

**Supplemental figure 2.**
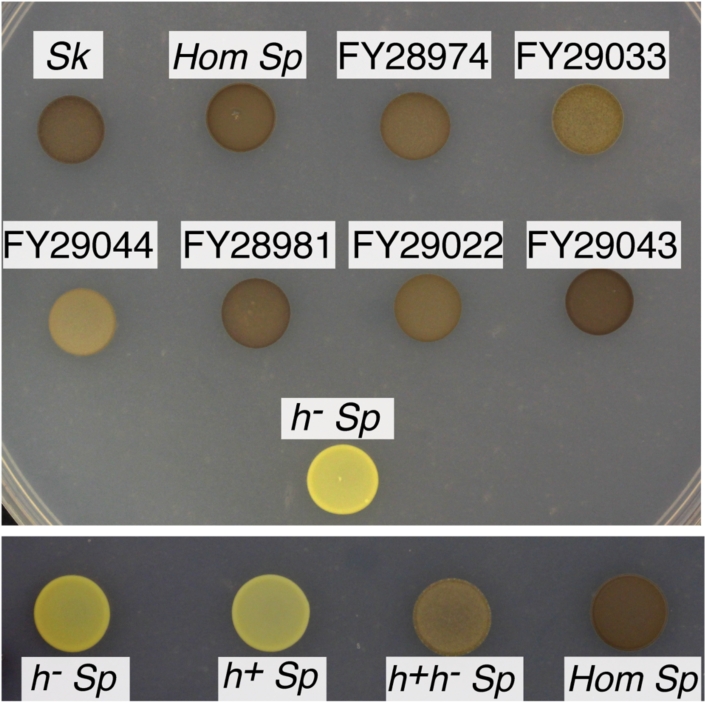
Homothallism of natural isolates. Saturated cultures were spotted on SPA+S to induce mating and meiosis. After five days at 25°C, cells were exposed to iodine vapor, which stains spores brown. The top panel shows natural isolates of homothallic (*Hom*) strains and *h- Sp* as a negative control. The bottom panel shows individual heterothallic strains, mixed complimentary heterothallic strains and a homothallic isolate of *Sp*.

**Supplemental figure 3.**
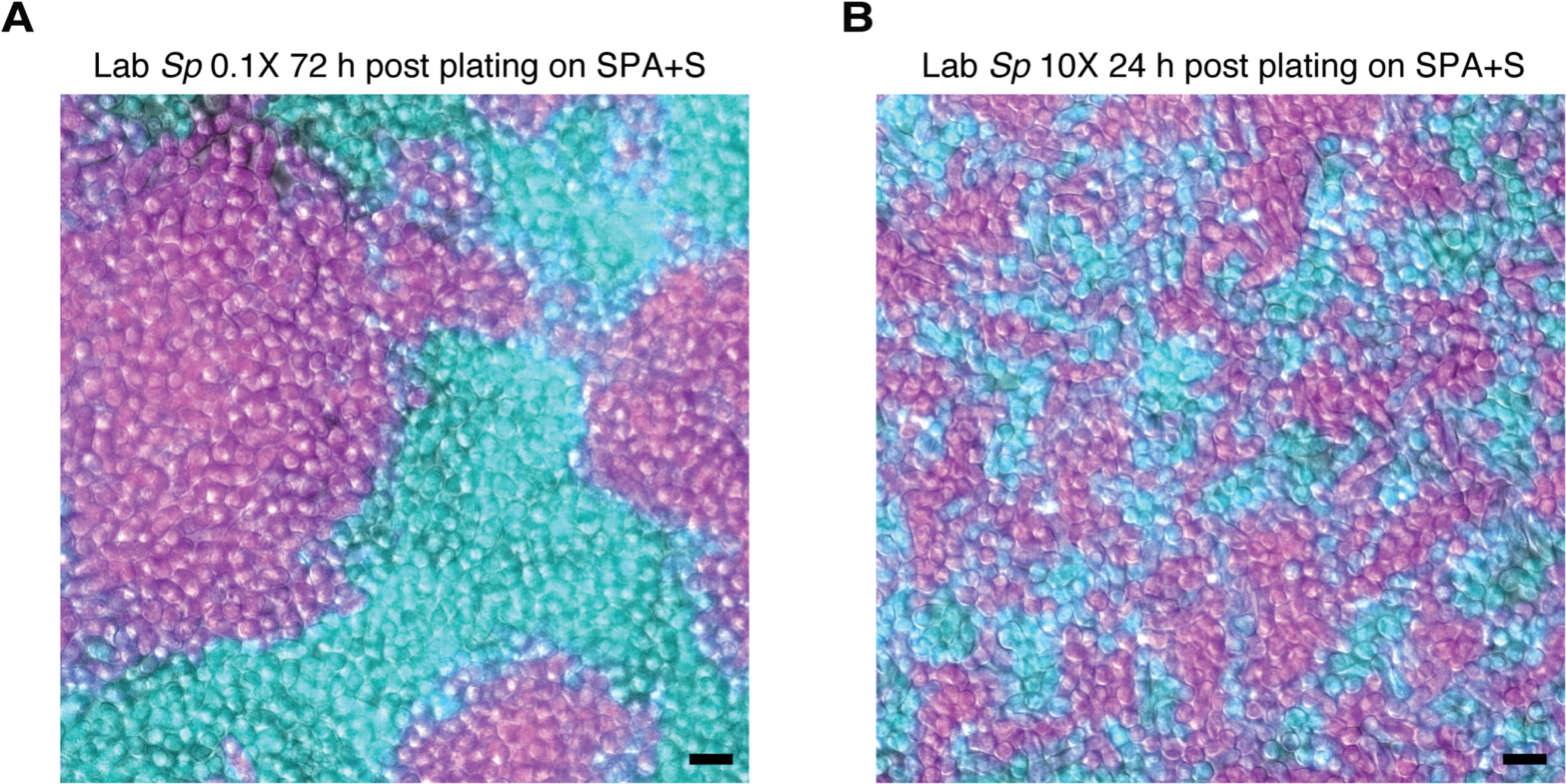
Density variation in plated cells. Homothallic GFP (cyan)- and mCherry (magenta)-expressing cells were mixed, plated on SPA+S, incubated at 25°C and then imaged. On this medium, cells tend to divide until confluent prior to mating. When plated at low density (0.1X; **A**), cells form large clusters of clonal cells after ∼72 hours. Cells mixed and plated at high density (10X; **B**) reach confluency after just 24 hours and form clusters with more contact zones between cells of different genotypes. Scale bars represents 10 µm.

**Supplemental figure 4.**
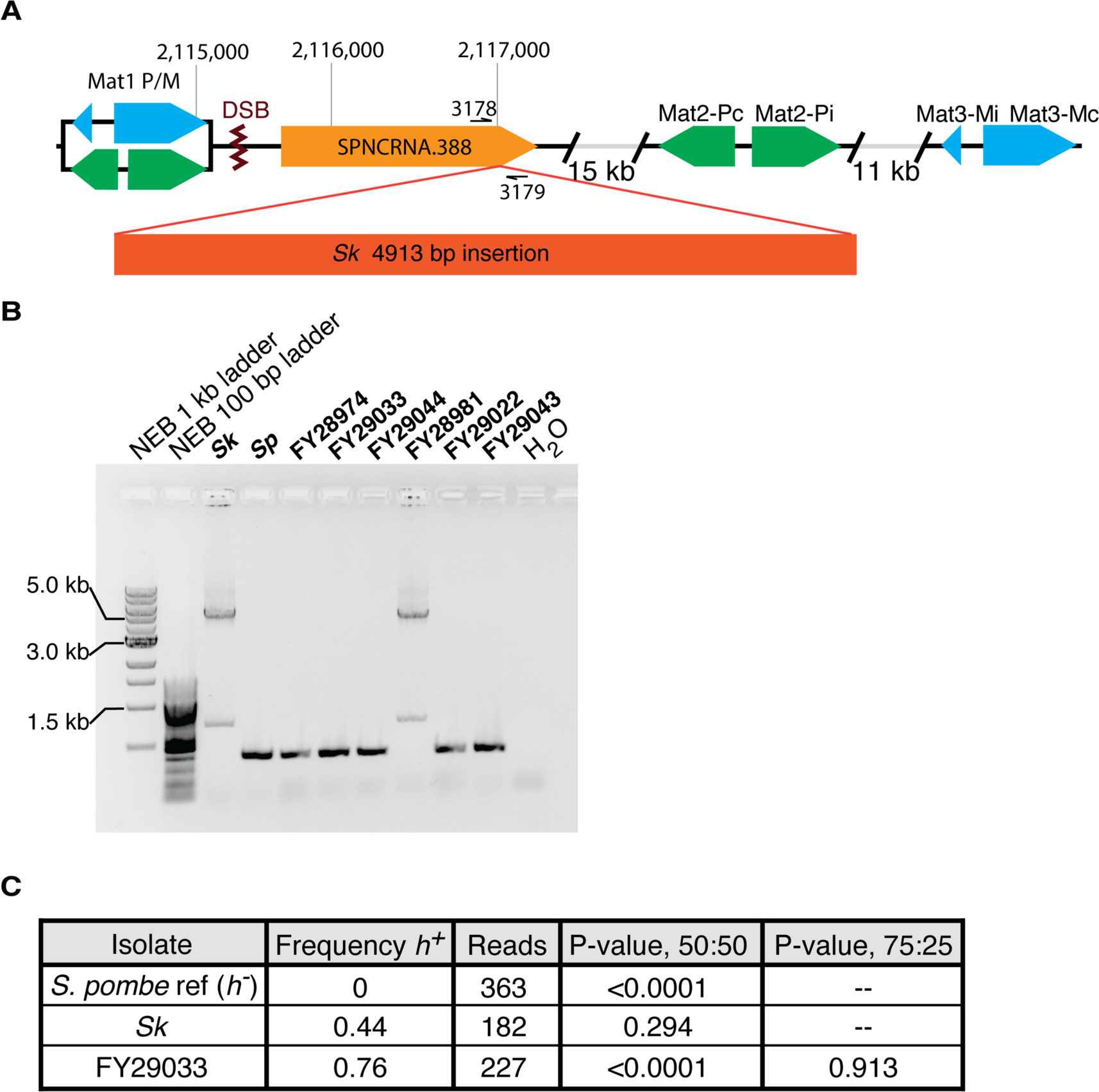
Mating-type locus variation in *S. pombe* isolates. **A)** Schematic representation of the mating-type locus. We found a 5 kb insertion of transposon sequences in the *Sk* isolate using published Illumina mate-pair sequencing data (Eickbush et al. 2019). The insert is within SPNCRNA.388, near the double-strand break site (DSB) where switching initiates. **B)** PCR amplification of the 5 kb insertion using the oligos 3178 and 3179 shown in **A** suggests the insertion is also in the FY28981 isolate. **C)** We aligned long Nanopore sequencing from the indicated strains to the *Sp* reference genome *mat* locus, which is *h-*. We then quantified the frequency of *h+* alleles from the aligned reads. We used Fisher’s exact tests to compare our observations to the indicated ratios.

**Supplemental figure 5.**
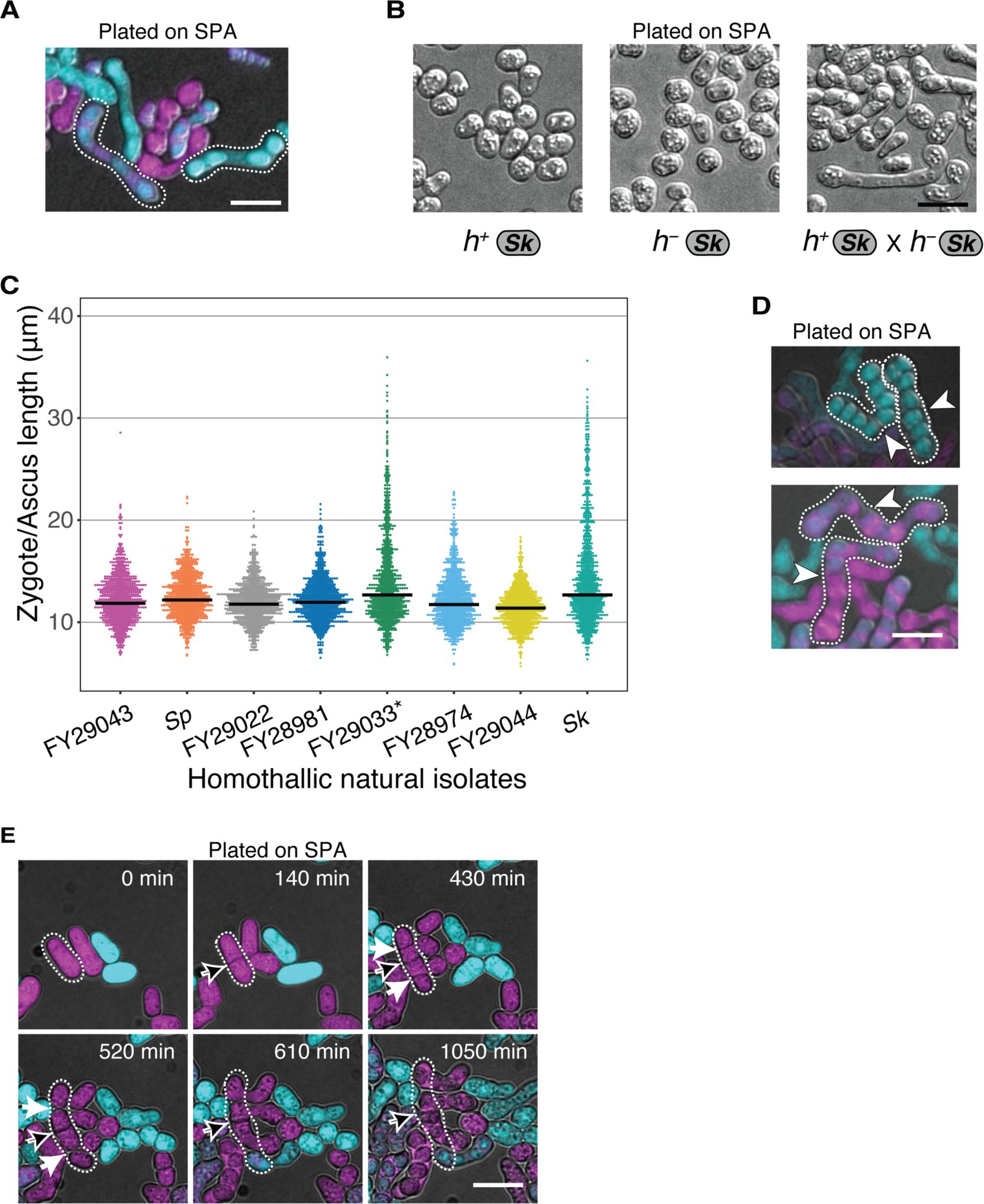
Variation in cell division and asci phenotypes in natural isolates. **A)** Examples of *Sk* long asci formed by GFP- and mCherry-expressing cells mated on SPA. **B)** *Sk* cells of each mating type were plated to SPA plates individually or in combination and imaged after 48 hours at 25°C. Cells only form long projections (shmoos) in response to cells of the opposite mating type. **C)** Tip-to-tip length (tracing through the center of the cells) of zygotes and asci of distinct natural isolates. Cells were plated on SPA and incubated for 24 hours at 25°C. *FY29033 cells showed a clumpy phenotype upon nitrogen starvation. **D)** Examples of *Sk* asci fused at uncleaved septa. White arrows indicate uncleaved septa. **E)** Time-lapse showing *Sk* cells with a septum (black arrow) that remained uncleaved after two rounds of fission. Additional septa (white arrows) are also shown that cleaved prior to mating. Scale bar represents 10 µm.

**Supplemental figure 6.**
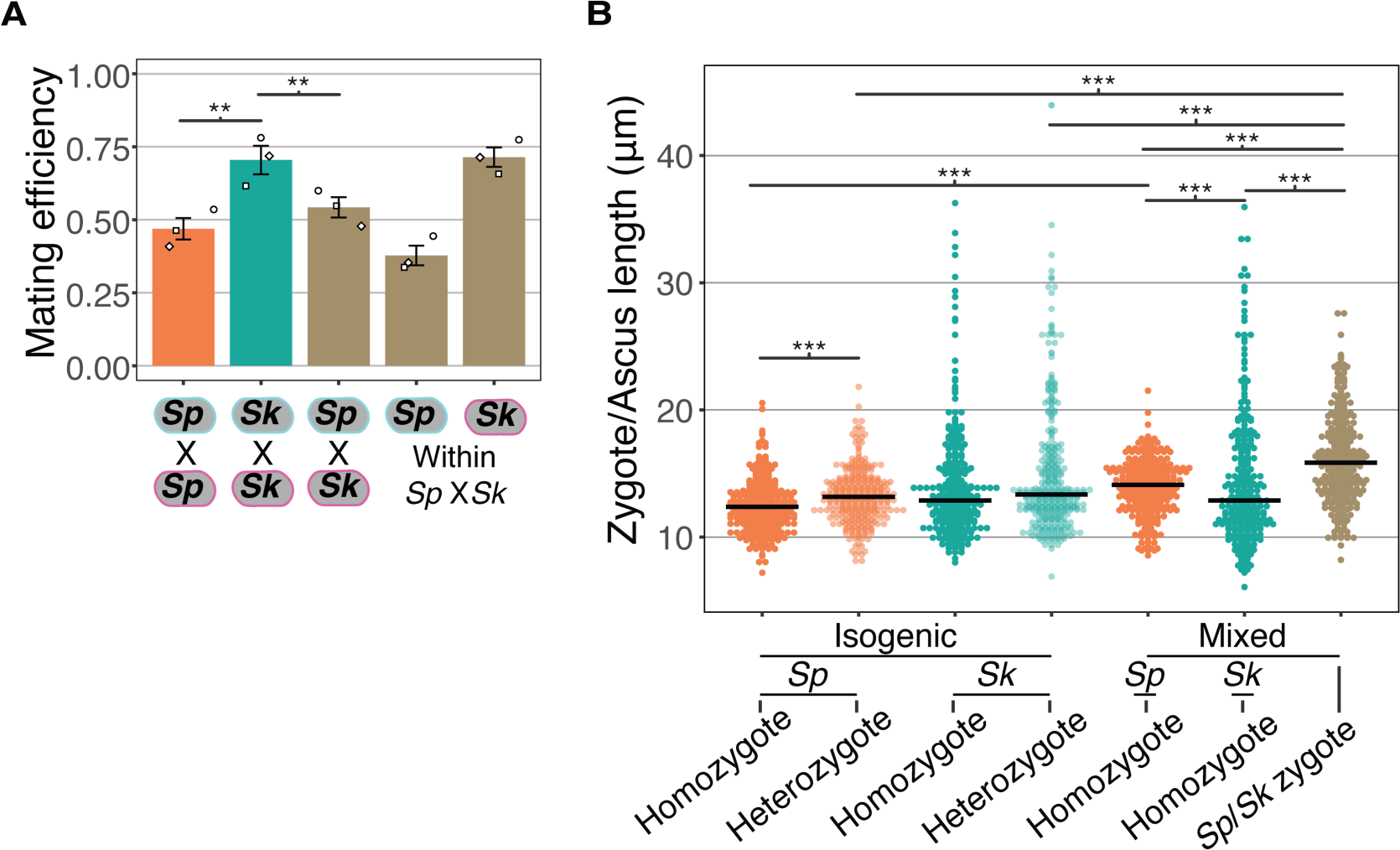
Mating efficiency and asci length of *Sp/Sk* heterozygotes. **A)** Mating efficiency calculated from still images taken after 24 hours on SPA at 25°C from isogenic and mixed isolate crosses. The open shapes represent values from three biological replicates per cross. ** indicates p-value <0.01, t-test. **B)** Length of zygotes/asci from isogenic and mixed isolate crosses of *Sp* and *Sk*. Cells were plated on SPA at 1X density and incubated for 24 hours at 25°C before being imaged. For each category (e.g., homozygote) of zygote, 300 cells were sampled from three independent biological replicates. Multiple Wilcoxon Rank Sum Tests, Bonferroni corrected; ***, p-value<0.005.

**Supplemental figure 7.**
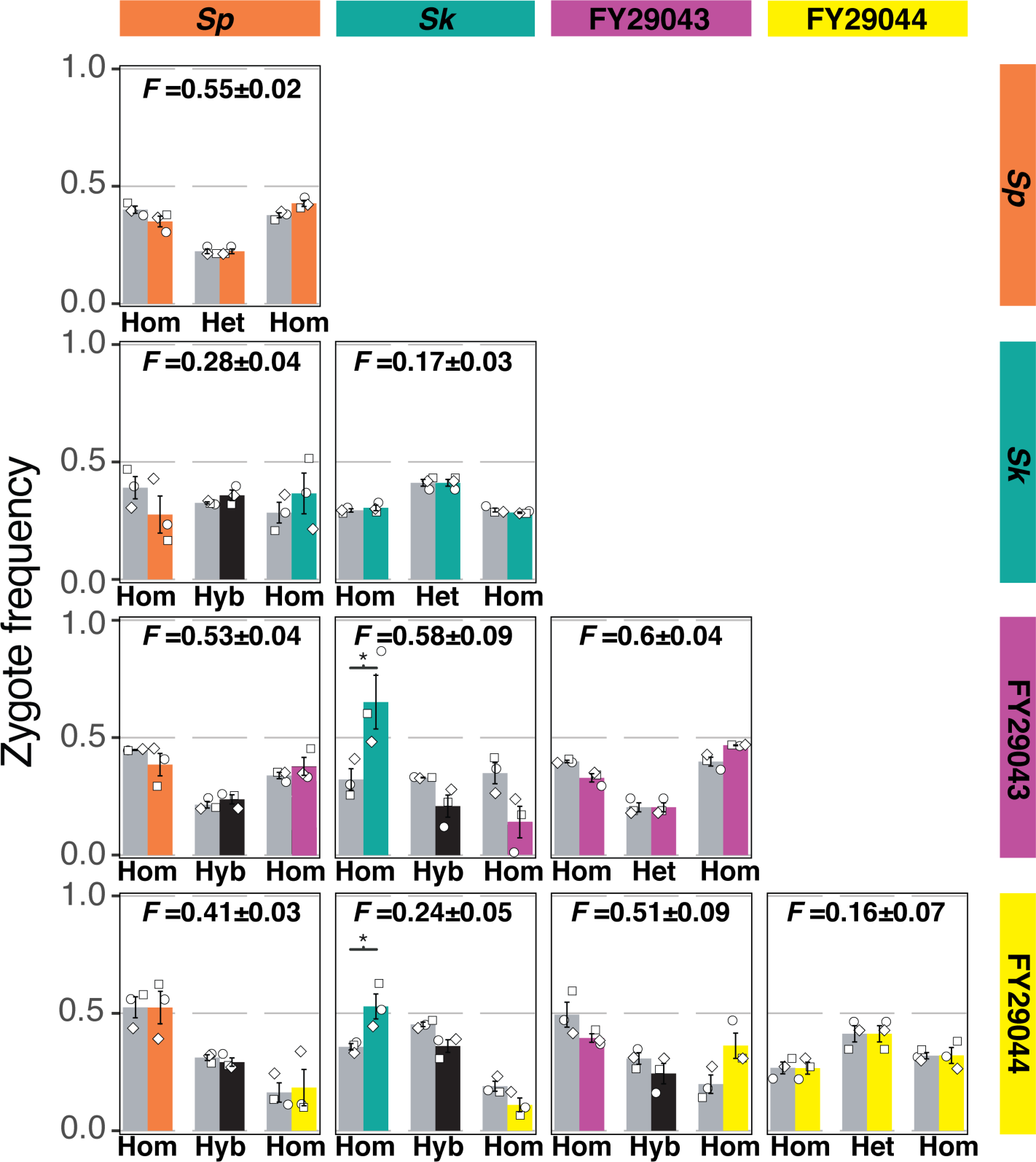
Mostly additive inbreeding phenotypes in crosses between natural isolates. Four natural isolates with distinct mating efficiencies and inbreeding coefficients were mixed and mated in all pair-wise combinations. After 24 hours on SPA at 25°C, cells were imaged and the mating efficiencies and inbreeding coefficients were calculated using fluorescent markers scored from still images, obtained as in Figure 1A. On the outer diagonal are isogenic crosses color-coded with the respective natural isolate. For the non-isogenic crosses, the observed homozygotes (hom) are color-coded by the respective parent isolate. The observed frequencies of hybrid (hyb) zygotes and asci are plotted in black. The expected values (gray bars) account for the inbreeding coefficient and mating efficiency from each isolate assuming the parental phenotypes are additive. Inbreeding coefficients (F values) and standard error (+/-) for each cross are indicated above each plot. The open shapes represent experimental replicates. * indicates p-value<0.05, one-tailed t-test.

**Supplemental figure 8.**
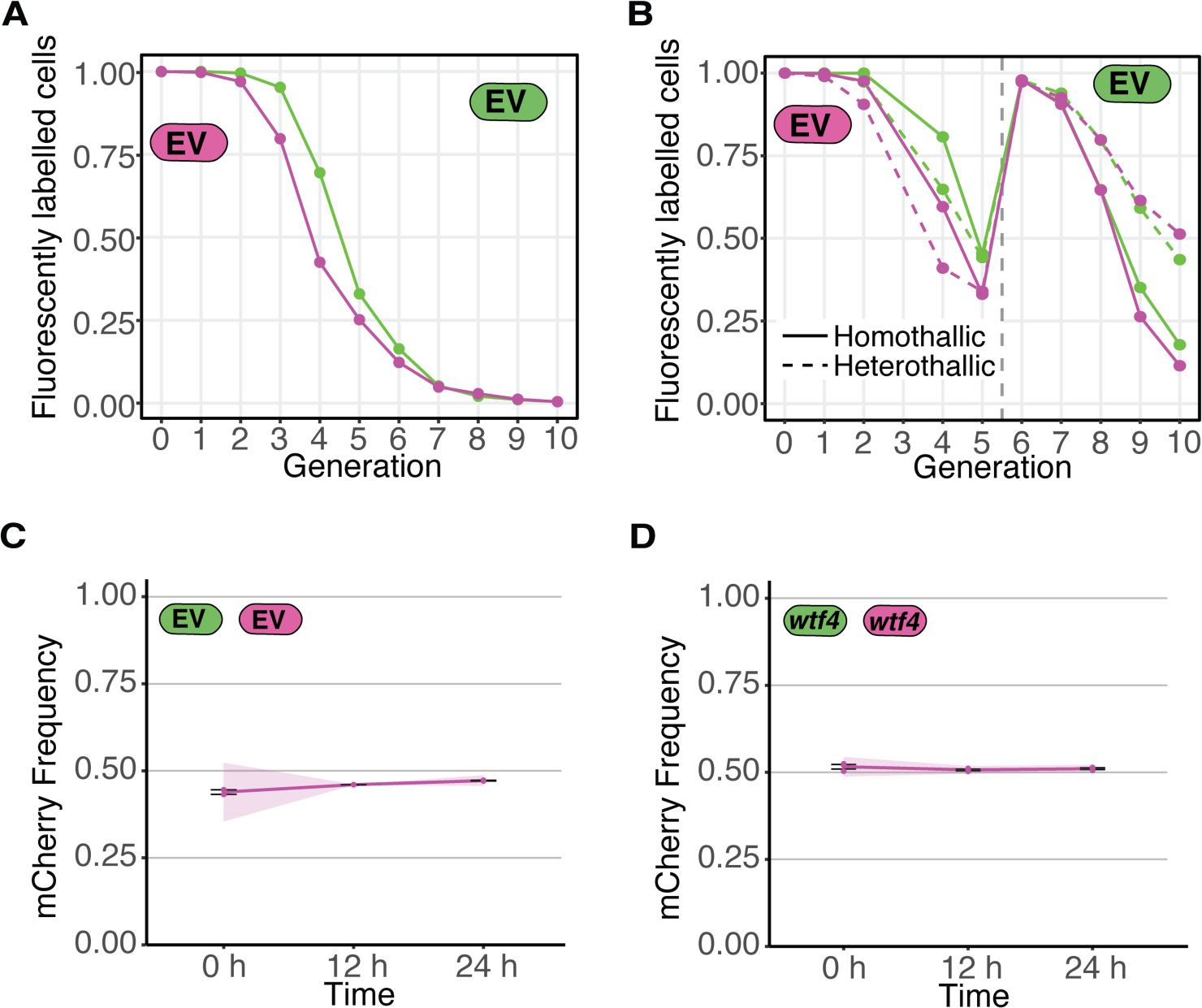
Loss of fluorescent markers in experimental evolution analyses and vegetative growth competition between fluorescently labelled cells. A) Fluorescent marker loss from cell populations carrying only one type of fluorescent marker (GFP or mCherry) linked to an empty vector (EV). Cells were grown as in Figure 4A in parallel to the cell populations depicted in Figure 4 B-C. B) Similar to A, but both heterothallic and homothallic cells were analyzed in parallel to the cell populations depicted Figure 4 D-E. The cells were sorted after generation 5 to remove non-fluorescent cells (vertical grey dotted line). C-D) We sorted GFP- and mCherry-positive cells from the progeny of a single replicate after nine meiotic generations. Sorted cells were taken from control crosses that are homozygous for *wtf4* locus (Supplemental figure 9 C-D). We then competed an equal ratio of GFP and mCherry cells starting with 30 µL of each strain into 600 µL of fresh YEL media for 12 hours at 32°C. Then 60 µL of culture was transferred to 600 µL fresh YEL media for 12 hours of growth C) Competition cultures during vegetative growth of strains with empty vector. D) Competition cultures during vegetative growth of strains carrying *wtf4*.

**Supplemental figure 9.**
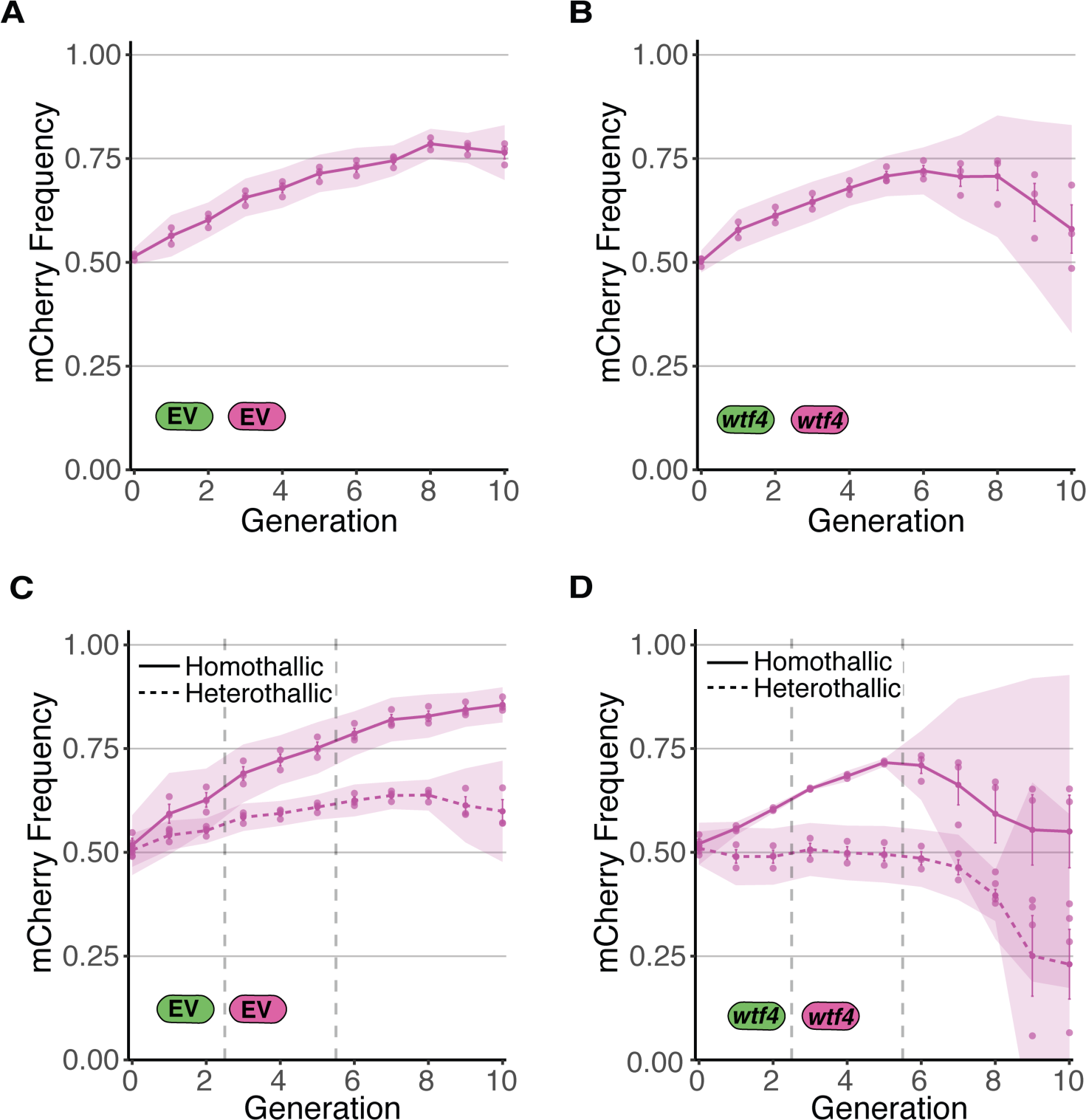
Allele transmission in homozygous control homothallic and heterothallic populations. **A)** Homothallic population with both mCherry and GFP marking EV alleles. The individual spots represent experimental replicates and the shaded areas around the lines represent 95% confidence intervals. **C)** Same experimental setup as in **B**, but with both markers linked to a *wtf4* allele. **C-D)** Repeats of the experiments shown in **B** and **C** with two alterations. First, these experiments tracked heterothallic populations (dotted lines) in addition to homothallic cells (solid lines). Second, the populations were sorted by cytometry at generations 3 and 5 (vertical long-dashed lines) to remove non-fluorescent cells.

